# Oligodendrocytes may provide the extra density of α-synuclein filaments from multiple system atrophy

**DOI:** 10.64898/2026.10.29.762735

**Authors:** Dongqi Zhang, Magdalena Matic, Airi Tarutani, Alexey G. Murzin, Melissa Huang, Jennifer A. Macdonald, Taxiarchis Katsinelos, Patrick C. Cullinane, Kazuko Hasegawa, Zane Jaunmuktane, Masato Hasegawa, Nadia Stefanova, Michel Goedert, Sjors H.W. Scheres

**Author notes:** These authors jointly supervised this work (Michel Goedert, Sjors H.W. Scheres).

## Abstract

Prion-like templated seeding of α-synuclein filaments is thought to underlie the spreading of multiple system atrophy (MSA) and other synucleinopathies through the brain. Here, we used α-synuclein filaments extracted from the brains of individuals with MSA to induce seeded aggregation in undifferentiated human neuroblastoma SH-SY5Y cells, human iNeurons, mixed primary murine cortical cells and murine oligodendrocyte-enriched primary cultures, all overexpressing human α-synuclein, followed by determination of the cryo-EM structures of the seeded filaments. Whereas MSA seeds comprise two protofilaments with an unidentified non-proteinaceous density located between protofilaments, seeding resulted in the formation of filaments with a single protofilament. Only the seeded structures derived from oligodendrocytes contained an extra density that resembled that of the seeds. This suggests that oligodendrocytes may provide the chemical component that is responsible for the extra density at the protofilament interface of MSA filaments. In support, the injection of MSA seeds into the striatum of transgenic mice expressing human α-synuclein under the control of the oligodendrocyte-specific proteolipid protein promoter, also yielded seeded filaments with a single protofilament and a similar extra density.

## Introduction

Filamentous inclusions of α-synuclein are a hallmark of several neurodegenerative diseases, collectively termed synucleinopathies. They include Parkinson’s disease (PD), dementia with Lewy bodies (DLB) and multiple system atrophy (MSA), as well as several rarer diseases^1^. Dominantly inherited missense and gene dosage mutations in *SNCA*, the α-synuclein gene, cause familial forms of PD and DLB with abundant Lewy pathology, supporting the notion that α-synuclein filaments play a central role in disease^2,3^. Moreover, sequence variation in the regulatory region of *SNCA* is associated with increased expression of α-synuclein in idiopathic PD^4^. α-Synuclein filaments are amyloids, small amounts of which can induce the assembly of monomeric protein into more filaments by templated seeding. Thereby, α-synuclein filaments provide the molecular basis for the prion-like mechanisms by which synucleinopathies are thought to spread through the brain.

MSA is a sporadic condition that is less common, but more aggressive than PD and DLB, with disease progression taking place over a period of 5-6 years and death occurring within 10 years of disease onset^5^. Parkinsonian (MSA-P) and cerebellar (MSA-C) subtypes are well established^6^. MSA-P is characterised by degeneration of the striatonigral system, giving rise to parkinsonian features that respond only poorly to levodopa. The olivopontocerebellar system degenerates in MSA-C, resulting in ataxia and gait abnormalities. Autonomic failure is characteristic of both MSA-P and MSA-C.

Irrespective of subtype, the predominant neuropathological characteristics of MSA are abundant filamentous glial cytoplasmic inclusions in oligodendrocytes (GCIs, also known as Papp-Lantos bodies)^7^; the number of GCIs correlates with the severity of neurodegeneration in specific brain regions in both MSA-P and MSA-C^8^. GCI filaments are made of α-synuclein phosphorylated at serine 129 (pS129)^9–12^. Filamentous neuronal cytoplasmic inclusions of α-synuclein are also present in MSA, in particular in the brainstem^13^. pS129 α-synuclein-positive protein can be found in cells without inclusions^12,14^. It has been suggested that pS129 α-synuclein-positive cells are particularly prone to inclusion formation. On the other hand, it has also been reported that pS129 α-synuclein protects against inclusion formation^15–17^.

Determination of the structures of amyloid filaments from human brains by electron cryo-microscopy (cryo-EM) has revealed that specific amyloid folds characterise different neurodegenerative diseases^18^; thus, the atomic structures of α-synuclein filaments extracted from the brains of individuals with MSA^19,20^ differ from those extracted from the brains of individuals with Lewy pathology (PD and DLB)^21,22^. Whereas α-synuclein filaments from PD and DLB brains consist of a single amyloid protofilament, i.e. with a single α-synuclein molecule on each layer of the helical filament, α-synuclein filaments from MSA brains consist of two types of filaments, each made of two different protofilaments. The proportions of type I and type II filaments vary between cases, but there is no evidence to suggest that MSA-P and MSA-C are characterised by distinct α-synuclein strains^19,23^. The same is true of neuronal and glial α-synuclein inclusions^19^.

Filaments with the Lewy fold and those extracted from the brains of individuals with MSA are characterised by the presence of extra densities of unknown composition that cannot be attributed to α-synuclein. In the Lewy fold, an extra density is situated within a pocket on the periphery of the ordered core that is coordinated by K32, K34, K43 and K45, as well as by Y39; in the MSA filaments, the extra density is at the interface between the two protofilaments in a pocket that is coordinated by K43, K45 and H50 from both protofilaments.

Templated seeding in the brain is believed to result in the formation of seeded filaments with the same structures as those of the seeds. To study the molecular mechanisms by which templated seeding leads to disease-specific filaments, it is necessary to develop model systems for seeded aggregation that reconstitute these filaments in a laboratory setting. However, this has so far not been possible for α-synuclein. Seeded assembly of purified recombinant α-synuclein into filaments has been observed to lead to filament structures that are different from those of the seeds^24,25^, and the same is also true of the intracerebral injection of preformed filaments of recombinant α-synuclein into the brains of transgenic mice heterozygous for human mutant A53T α-synuclein^26^.

In this study, we explored seeded aggregation in several cell culture models through the addition of seeds extracted from human brains with MSA type II α-synuclein filaments. To facilitate seeding within the time frame in which these cells are viable, we overexpressed α-synuclein constructs, with or without tags or mutations, using different expression strategies including transient transfection and viral transduction. For each cell culture model, we extracted seeded α-synuclein filaments using sarkosyl and determined their atomic structures by cryo-EM. All seeded α-synuclein filaments comprised only a single protofilament. When α-synuclein was overexpressed in oligodendrocytes, we did observe a density between K43 and K45 that was reminiscent of the density located at the interface of the two protofilaments in the MSA folds. In MSA brains, α-synuclein inclusions are typically observed in the cytoplasm of oligodendroglia^6^. Therefore, our results suggest that oligodendrocytes may provide the chemical compound(s) that are responsible for this extra density in the MSA folds. Previous studies described the generation of transgenic mouse lines overexpressing human α-synuclein in oligodendrocytes under the control of the proteolipid protein (PLP)^27,28^ promoter or the 2’3’-cyclic nucleotide 3-phosphodiesterase promoter^29^. Here we show that the intrastriatal injection of MSA seeds into PLP-αSyn transgenic mice also yielded seeded α-synuclein filaments with a single protofilament and a similar extra density.

## Results

### Structures of Triton X-100-extracted α-synuclein filaments from MSA brain

Sarkosyl is an anionic detergent that is being widely used for the extraction of amyloid filaments from human brains for cryo-EM studies^30,31^. Its ionic properties give it a strong ability to solubilise many components of brain tissues, while amyloid filaments retain their insoluble nature. Sarkosyl extraction of tau filaments from the brains of individuals with Alzheimer’s disease yielded the same cryo-EM structures as those of filaments obtained using immunopurification or water-based extraction^31–33^. Moreover, the same filament structures were also found in the brains of individuals with Alzheimer’s disease by electron cryo-tomography^34^. However, it is not known if the use of sarkosyl affects the structures of α-synuclein filaments extracted from human brains.

We therefore used Triton X-100, a non-ionic detergent, to extract α-synuclein filaments from the frontal cortex of an individual with FTLD-synuclein for whom we previously reported the cryo-EM structures of sarkosyl-extracted α-synuclein filaments^35^. Compared with sarkosyl, which has a negatively charged sarcosine-derived hydrophilic head group, Triton X-100 contains a hydrophilic polyethylene oxide chain, making it a milder detergent for protein solubilisation^36^. The resulting cryo-EM structures (Figure 1A,B), which were determined to a resolution of 3.0 Å, were identical to those obtained using sarkosyl extraction, as confirmed by an amyloid packing difference (APD)^37^ of 4.5% (Figure S1A) and an all non-hydrogen atom root mean square deviation of 0.36 Å. Position and shape of the unidentified density at the protofilament interface were very similar in both reconstructions (Figure 1B). We conclude that sarkosyl did not introduce artefacts in the cryo-EM structures of brain-derived MSA filaments. It is therefore also unlikely that the unidentified density corresponds to the detergents used for filament extraction. In the remainder of this study, we used sarkosyl to extract α-synuclein filaments from cell cultures and brain tissues.

**Figure 1:**
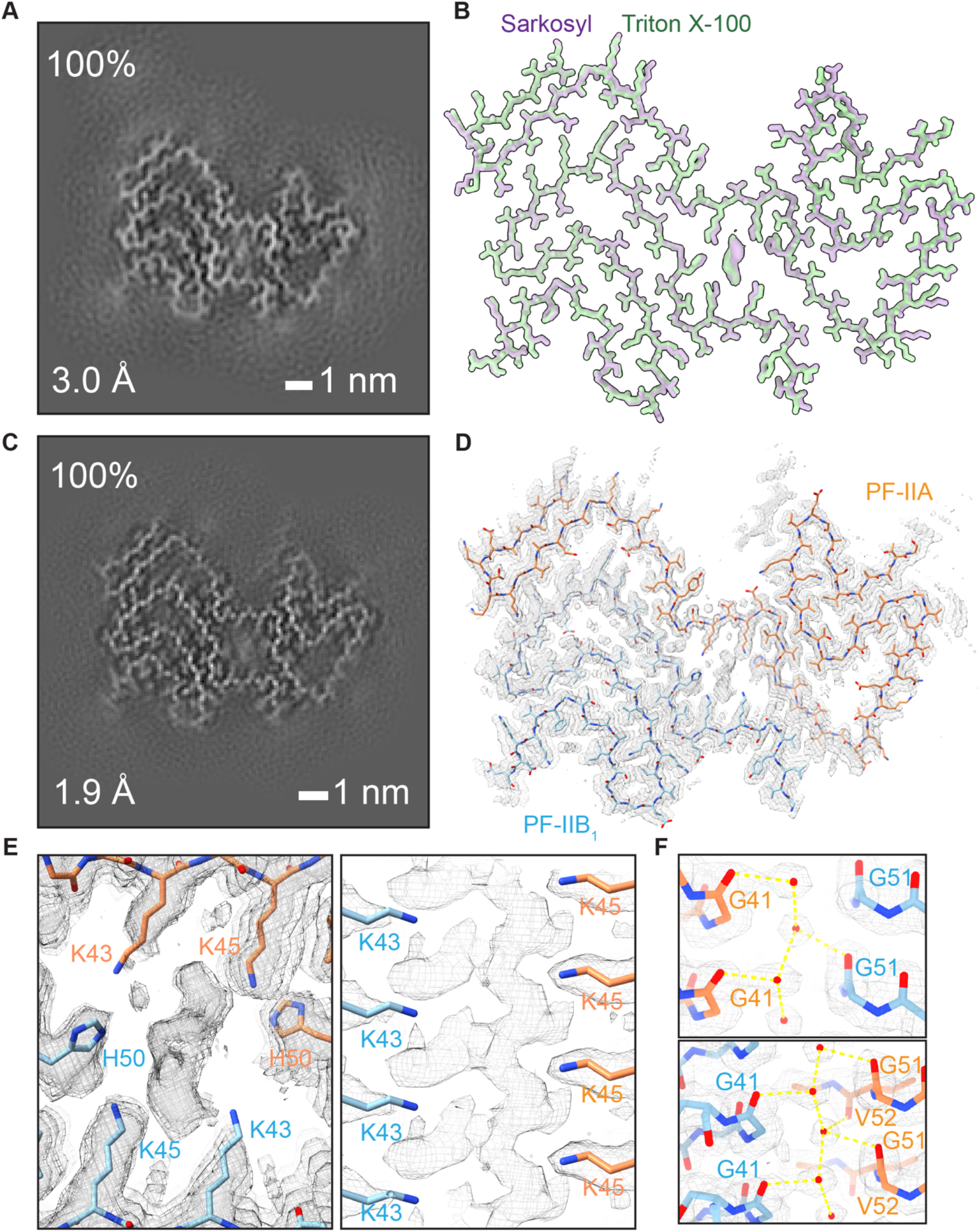
Cryo-EM structures of α-synuclein filaments from MSA brain. A. Cryo-EM cross-section of Triton X-100-extracted α-synuclein filament from MSA brain. The view is perpendicular to the helical axis, with a projected thickness of approximately one rung. The resolution of the reconstruction and the filament percentage are indicated. B. Backbone representation of the overlay of α-synuclein filaments from MSA brains extracted using either Triton X-100 or sarkosyl (PDB 9CX6). C. Cryo-EM cross-section of sarkosyl-extracted α-synuclein filament from MSA brain at 1.9 Å resolution. The view is perpendicular to the helical axis, with a projected thickness of approximately one rung. The resolution of the reconstruction and the filament percentage are indicated. D. Cryo-EM density map and atomic model of the 1.9 Å structure. E. Top and side views of the co-factor density in the 1.9 Å structure. F. Water densities at the interface between the two protofilaments of the 1.9 Å structure and corresponding hydrogen-bonding interactions.

### Structures of α-synuclein filaments from MSA brain at 1.9 Å resolution

We next set out to obtain more information about the extra density at the protofilament interface of MSA filaments. In an attempt to increase the resolution, we acquired 16,610 micrograph movies of sarkosyl-extracted α-synuclein filaments from the case with FTLD-synuclein^35^ using a G4 Titan Krios microscope with a Selectris-X energy filter and a Falcon4i detector^38^.

The reconstruction of 224,301 helical segments in RELION-5.1^39^ yielded a 1.9 Å resolution structure, in which densities for almost all non-hydrogen atoms of α-synuclein were resolved (Figure 1C-E). The unidentified density had an elongated tri-lobed shape with approximate dimensions in the XY-plane of 4x10 Å, partial per-layer separation in the Z-direction, and an approximate volume of 90 Å^3^ per layer. It did not connect to any of the four surrounding lysines. However, the resolution of the density did not improve as much as that of α-synuclein, suggesting that the molecules responsible are heterogeneous and/or do not obey the imposed helical symmetry, which makes it difficult to deduce insights into the density’s chemical composition.

The 1.9 Å reconstruction map revealed nine ordered water molecules inside the MSA filaments. Two pairs of ordered water molecules, on either side of the unidentified density, contributed to the interactions at the protofilament interface. On one side, two water molecules formed hydrogens bonds with the carbonyl groups of G51 and V52 of protofilament B and the carbonyl group of G41 of protofilament A; on the other side, they formed hydrogen bonds with the carbonyl groups of G51 and V52 of protofilament A and the carbonyl group of G41 of protofilament B (Figure 1F). The side chain of K80 in protofilament A was pointing down and was not connected to the nearby peptide-like density of unknown identity, indicating that it was unlikely that the density corresponded to ubiquitin.

### Cryo-EM structures of MSA-seeded α-synuclein filaments from human neuroblastoma SH-SY5Y cells

We explored if we could reconstitute the structures of MSA filaments from the human brain in two-dimensional cell culture models upon overexpression of α-synuclein and the addition of MSA seeds. We first investigated the seeding of SH-SY5Y cells transiently overexpressing wildtype or A53T human α-synuclein, or wildtype murine α-synuclein, with α-synuclein seeds extracted from the putamen of MSA case 5 (Figure 2A)^19^.

**Figure 2:**
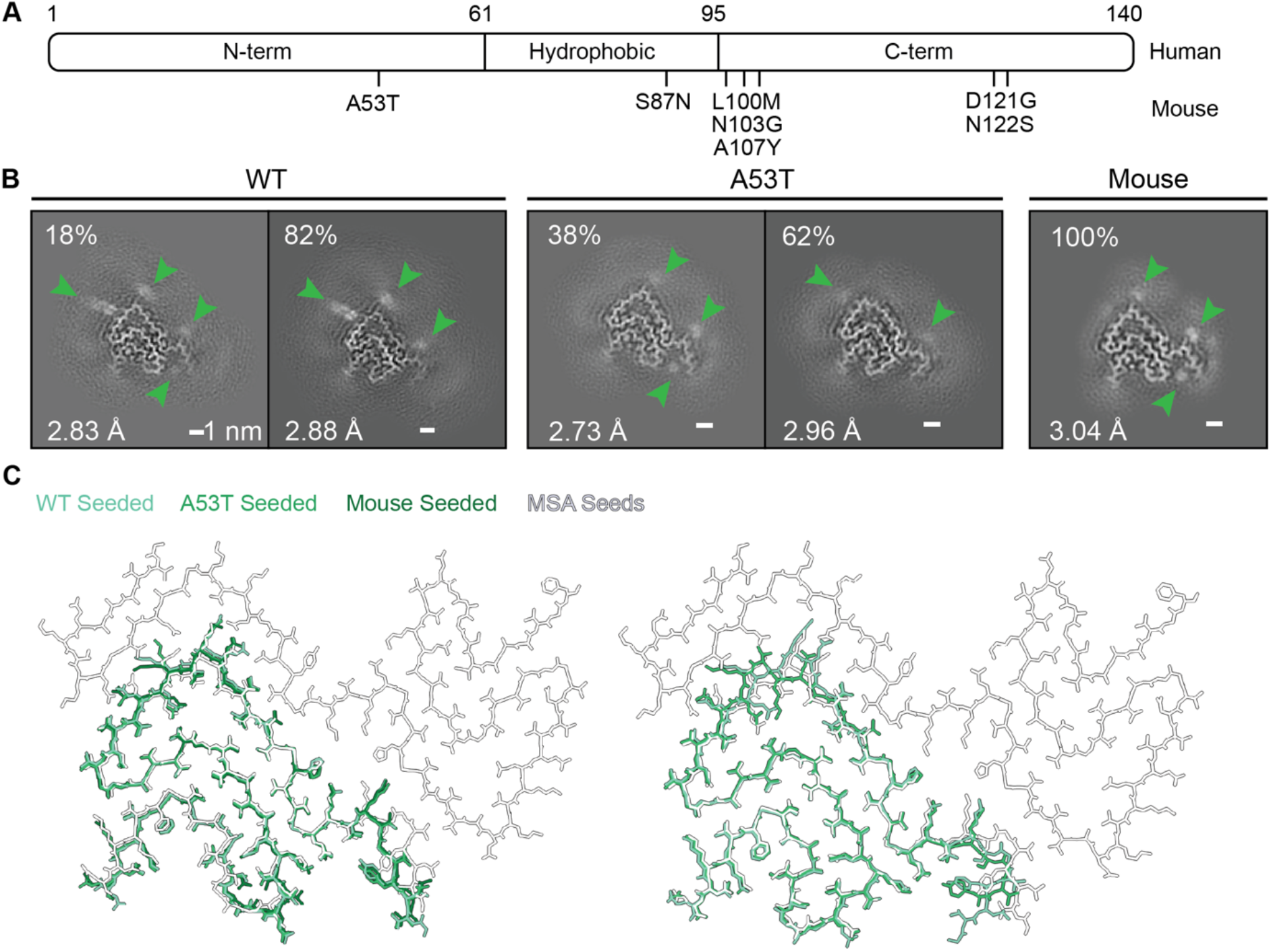
Cryo-EM structures of MSA-seeded α-synuclein filaments from human neuroblastoma SH-SY5Y cells. A. Sequence comparison between human and mouse α-synuclein. They differ by seven amino acids. B. Cryo-EM cross-sections of seeded α-synuclein filaments from SH-SY5Y cells overexpressing wildtype or A53T human and wildtype murine α-synuclein. C. Overlay of the atomic models of seeded filament structures with those of the MSA seeds (PDB 6XYP and 6XYQ).

Expression of wildtype or A53T human α-synuclein led to the formation of filaments with a single protofilament (singlets) in two distinct conformations that were nearly identical to the two conformations (PF-IIB_1_ and PF-IIB_2_) of protofilament IIB from the MSA seeds, with only minor conformational changes near the amino-terminal part of the ordered cores (Figure 2B,C). The (XY) APDs between the seeded structures and PF-IIB_1_ or PF-IIB_2_ of the seeds ranged from 3-16% (Figure S1B-C). When transiently transfecting wildtype murine α-synuclein, only singlets with the conformation of PF-IIB_1_ were observed. N87 (this residue is a serine in human α-synuclein), formed a hydrogen bond with T81, which stabilised the conformation of PF-IIB_1_ (Figure S1D). The XY-APD with the PF-IIB_1_ conformation of the seeds was 8% (Figure S1E).

None of the seeded filaments in SH-SY5Y cells recapitulated the structures of the MSA seeds, which comprise two non-identical protofilaments, and none of the maps showed extra densities between K43 and K45, which coordinate the unidentified density in MSA filaments. Other unidentified densities that surrounded the density for α-synuclein in the filaments from SH-SY5Y cells were distinct in the two different conformations (green arrows in Figure 2B). We conclude that SH-SY5Y cells do not provide the cellular environment necessary for the correct templated seeding of MSA filaments.

### Cryo-EM structures of MSA-seeded α-synuclein filaments from human iNeurons

We next explored seeded aggregation of α-synuclein in human pluripotent stem cell-derived neurons induced by the NGN2-OPTi-OX-hPSCs system^40^; we used lentiviruses to overexpress wildtype human α-synuclein three days after differentiation with doxycycline. We used two distinct carboxy-terminally tagged α-synuclein constructs: either a human influenza virus hemagglutinin (HA) tag or a combined Venus-HA tag. α-Synuclein filaments extracted from the frontal cortex of the individual with FTLD-synuclein^35^ were used for seeding transduced iNeurons four days after they had been induced to differentiate. These seeds are identical in structure to those from the putamen of typical MSA^35^. Following three more weeks in culture, immunohistochemistry and immuno-EM showed the formation of seeded α-synuclein filaments (Figures 3B, S2A). By cryo-EM, 2D class averages of the extracted filaments with the Venus-HA tag showed additional fuzzy densities, which we attribute to the Venus tag, when compared to 2D class averages of filaments that only contained an HA-tag (Figure S2B). Cryo-EM reconstructions of the Venus-HA-tagged and the HA-tagged seeded α-synuclein filaments showed singlets in similar conformations, indicating that the size of the carboxy-terminal tags did not influence the structures of the seeded filaments (Figure 3C).

**Figure 3:**
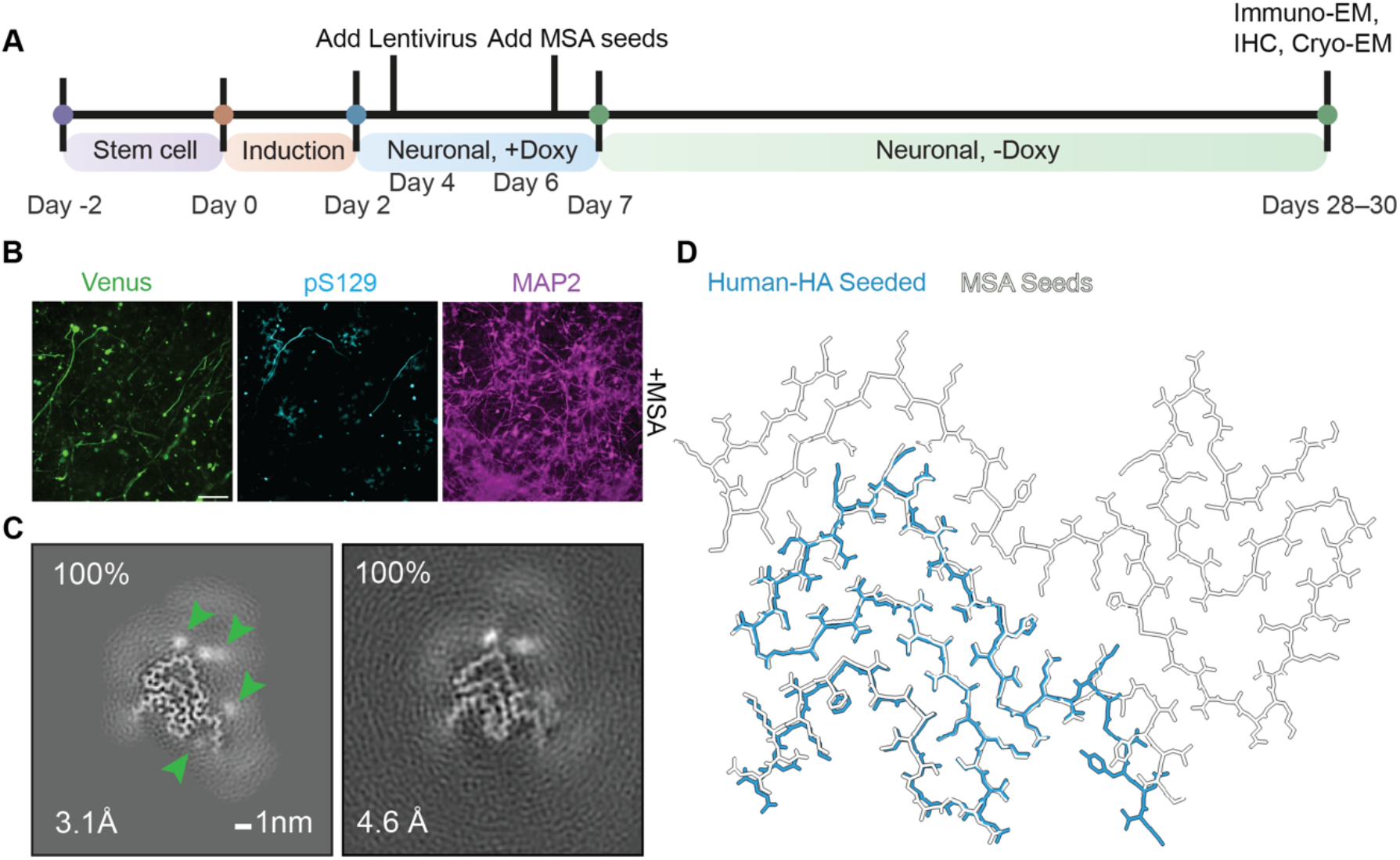
Cryo-EM structures of MSA-seeded α-synuclein filaments from human iNeurons. A. Schematic representation of experiments. B. Immunohistochemical staining of iNeurons for the Venus-tag, pS129 α-synuclein and MAP2 21-23 days after the addition of MSA seeds. Scale bar, 100 μm. C. Cryo-EM cross-sections of seeded α-synuclein filaments from iNeurons overexpressing α-synuclein-HA (left) or α-synuclein-Venus-HA (right). D. Overlay of the atomic models of the seeded α-synuclein-HA filament structures with those of the MSA seeds (1.9 Å resolution structure).

Despite the cellular environment being that of human nerve cells, cryo-EM reconstructions of the Venus-HA-tagged and the HA-tagged MSA-seeded α-synuclein filaments showed only singlets. The seeds were made of MSA type II α-synuclein filaments (PF-IIA/PF-IIB_1_)^35^, but we only observed singlets with the PF-IIB_1_ conformation for the seeded filaments (Figure 3D). Similar to the filaments seeded in SH-SY5Y cells, these filaments did not contain extra densities between K43 and K45, but they did show other extra densities similar to those in filaments seeded in SH-SY5Y cells (green arrows in Figure 3C). The XY-APD with the MSA PF-IIB_1_ conformation of the seed was 6% (Figure S1F).

### Cryo-EM structures of MSA-seeded α-synuclein filaments from mixed primary murine cortical cell cultures

Our next attempt to reconstitute MSA filaments in cell culture used mixed primary cortical cultures from P0/1 C57BL/6J-ola mice, which do not express α-synuclein because of a deletion of *SNCA*^41^. These cultures comprised a mixture of different cell types, which we reasoned could play a role in the formation of MSA filaments.

On day 0, we used an adeno-associated virus (AAV) to overexpress wildtype human α-synuclein with a carboxy-terminal Venus tag under the control of the neuronal synapsin 1 promoter. Four days later, we added MSA seeds from the FTLD-synuclein case. After another 12-14 days in culture, immunohistochemistry and immuno-EM showed the formation of seeded filaments inside neurons (Figures 4B and S2A). By cryo-EM, 2D class averages of the sarkosyl-extracted filaments showed fuzzy densities around the filaments because of the Venus tag (Figure S2B), and 3D reconstructions again revealed 100% singlets with the PF-IIB_1_ conformation (Figure 4C, D), with extra densities similar to those observed with SH-SY5Y cells and iNeurons. The XY-APD with the MSA PF-IIB_1_ of the seeds was 6% (Figure S1G).

**Figure 4:**
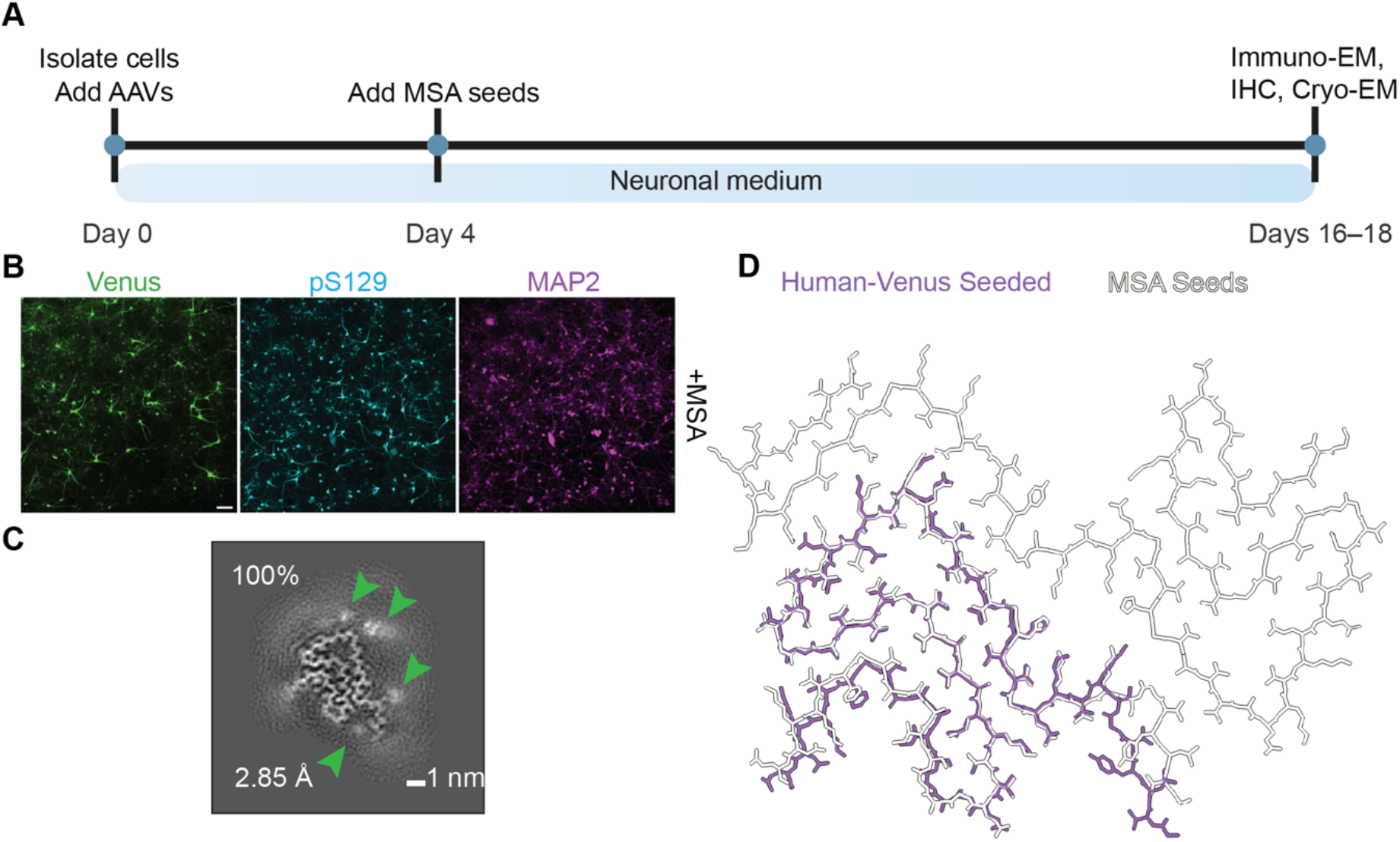
Cryo-EM structures of MSA-seeded α-synuclein filaments from mixed primary murine cortical cells. A. Schematic representation of experiments. B. Immunohistochemical staining of mixed primary murine cortical cells for the Venus-tag, pS129 α-synuclein and MAP2 12-14 days after the addition of MSA seeds. Scale bar, 100 μm. C. Cryo-EM cross-section of seeded α-synuclein filament from mixed primary murine cortical cells overexpressing α-synuclein-Venus. D. Overlay of the atomic models of the seeded α-synuclein-Venus filament structures with those of the MSA seeds (1.9 Å resolution structure).

### Cryo-EM structures of α-synuclein filaments from murine oligodendrocyte-enriched primary cultures

MSA is characterised by abundant GCIs in oligodendrocytes^7^. Because none of the cell culture models described so far reconstituted the structures of the seeds, we reasoned that the cellular environment of oligodendrocytes might be critical. To test this hypothesis, we developed an *in vitro* cell seeding model. Initial attempts to seed cultures of oligodendrocytes were unsuccessful, because they died during the two-week timespan of the seeding assays. We therefore co-cultured oligodendrocytes and neurons, which resulted in their survival (Figure 5A, S4).

**Figure 5:**
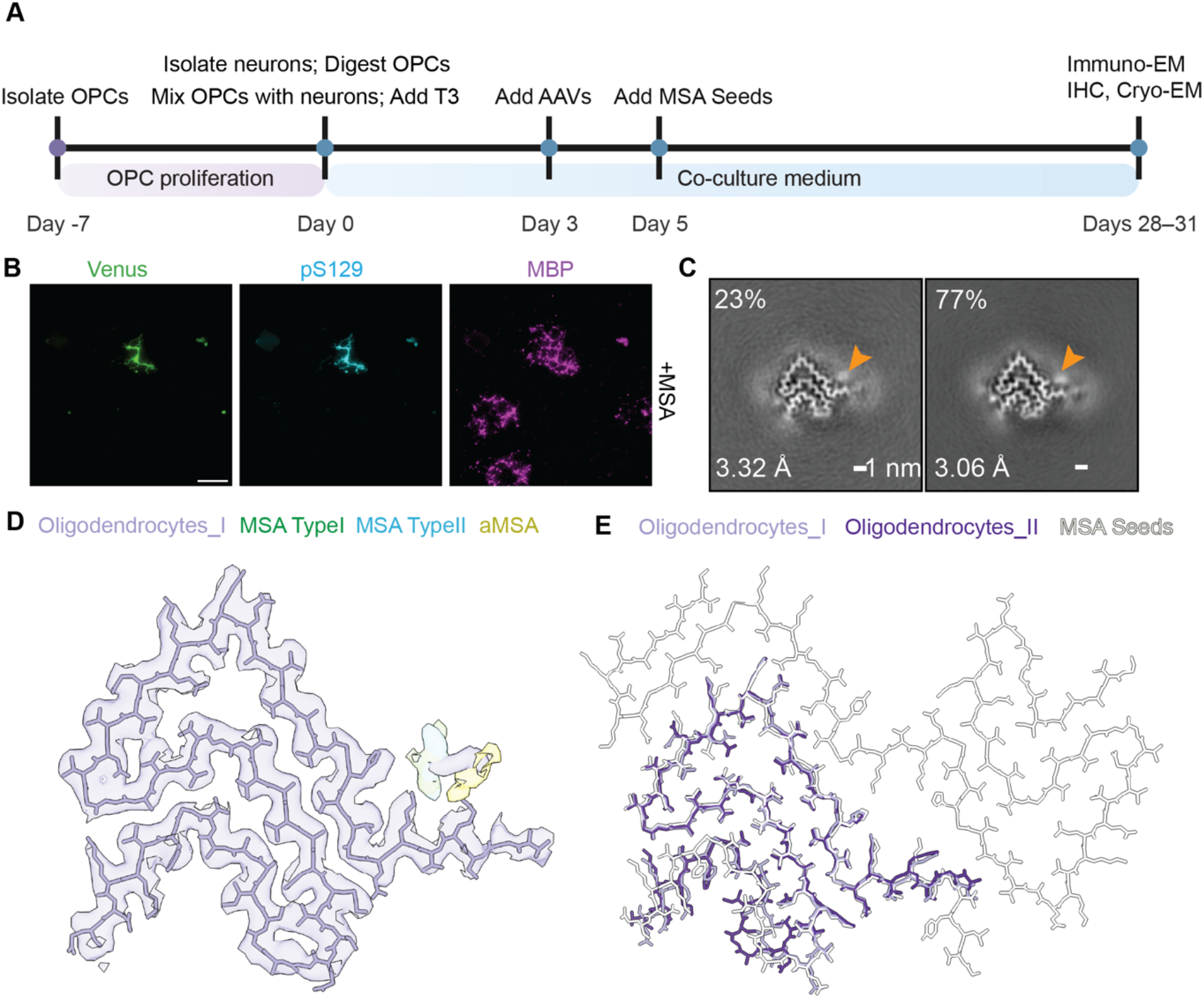
Cryo-EM structures of MSA-seeded α-synuclein filaments from murine oligodendrocyte-enriched primary cultures. A. Schematic representation of experiments. B. Immunohistochemical staining of oligodendrocyte-enriched primary cultures for the Venus-tag, pS129 α-synuclein and myelin basic protein (MBP) 23-26 days after the addition of MSA seeds. Scale bar, 100 μm. C. Cryo-EM cross-sections of seeded α-synuclein filaments from murine oligodendrocyte-enriched primary cultures overexpressing α-synuclein-Venus. Two different conformations of seeded structures from oligodendrocytes are observed: the left one adopts the PF-IIB_1_ conformation, while the right one adopts the PF-IIB_2_ conformation. D. Overlay of the non-proteinaceous densities in the cryo-EM maps of the MSA-seeded structures with those of known α-synuclein filament structures from the brains of individuals with MSA (MSA Type I, EMD-10650; MSA Type II_1_, EMD-45979; aMSA, EMD-70295). E. Overlay of the atomic models of the seeded α-synuclein-Venus filament structures with those of the MSA seeds (1.9 Å resolution structure).

We purified oligodendrocyte progenitor cells (OPCs) from the cerebral cortex of P0/1 mice by density gradient centrifugation^42^. Once the OPC colonies had reached confluence, they were digested by Accutase (3ml/flask) and mixed with primary cells from P0/1 mouse cortex; T3 thyroid hormone, a differentiation factor for oligodendrocytes^42^, was also added. Five days after establishing the co-cultures, we added an AAV with an Olig001 capsid^43^ to achieve oligodendrocyte-specific overexpression of Venus-tagged human α-synuclein under the control of the myelin basic protein promoter^44^. After another three days, we added MSA seeds from the FTLD-synuclein case^35^. Following a 23-26-day incubation with seeds, oligodendrocyte-specific seeding of expressed α-synuclein was shown by the co-localisation of α-synuclein inclusions and myelin basic protein (Figure 5B). Sarkosyl-extracted filaments from these cultures labelled with a GFP antibody by immuno-EM, and 2D class averages again showed fuzzy densities from the Venus tag (Figure S2B).

Three-dimensional reconstructions of the seeded filaments revealed singlets, but now with both PF-IIB_1_ and PF-IIB_2_ conformations (23 and 77%, respectively) (Figure 5C). It is unclear why we observed two conformations for these filaments, whereas the same seeds yielded only PF-IIB_1_ filaments in iNeurons and primary cortical cultures. The PF-IIB_1_ and PF-IIB_2_ conformations of the α-synuclein singlets were nearly identical to the corresponding structures obtained with the cellular models described above, with XY-APDs of 11% (Figure S1H,I). Importantly, both reconstructions now showed an extra density between K43 and K45, in a position similar to that of the unidentified density in MSA filaments (Figure 5D,E).

### Cryo-EM structures of MSA-seeded α-synuclein filaments from PLP-αSyn transgenic mice

We next injected α-synuclein filament seeds from the individual with FTLD-synuclein^35^ into the dorsal striatum of PLP-αSyn transgenic mice to determine the cryo-EM structures of the seeded filaments. This transgenic mouse line expresses human α-synuclein in oligodendrocytes under the control of the PLP promoter^27,28^. Using sarkosyl extraction, we did not observe filaments in non-injected PLP-αSyn mouse brains (Figure S5A). Accordingly, there was no staining with the amyloid dye pFTAA^45^ (Figure 6A), in agreement with previous findings using Thioflavin T^46^. However, seven months after the intracerebral injection of MSA seeds, transgenic mouse brains showed strong staining with pFTAA, especially in white matter tracts (*corpus callosum*, *capsula externa*, *capsula interna*, anterior commissure and *crus cerebri*) (Figure 6B). We also observed filaments in injected PLP-αSyn mouse brains (Figure S5B). Unlike pFTAA staining, there was substantial pS129 staining both without and with the injection of seeds (Figure S6A,B). Following MSA seed injection, pFTAA staining co-localised to a significant extent with that for pS129 (Figure S6C-E). As expected, following seed injection, pFTAA staining also co-localised with immunofluorescence for p25α, a marker for oligodendrocytes^47^ (Figure 6C-E).

**Figure 6:**
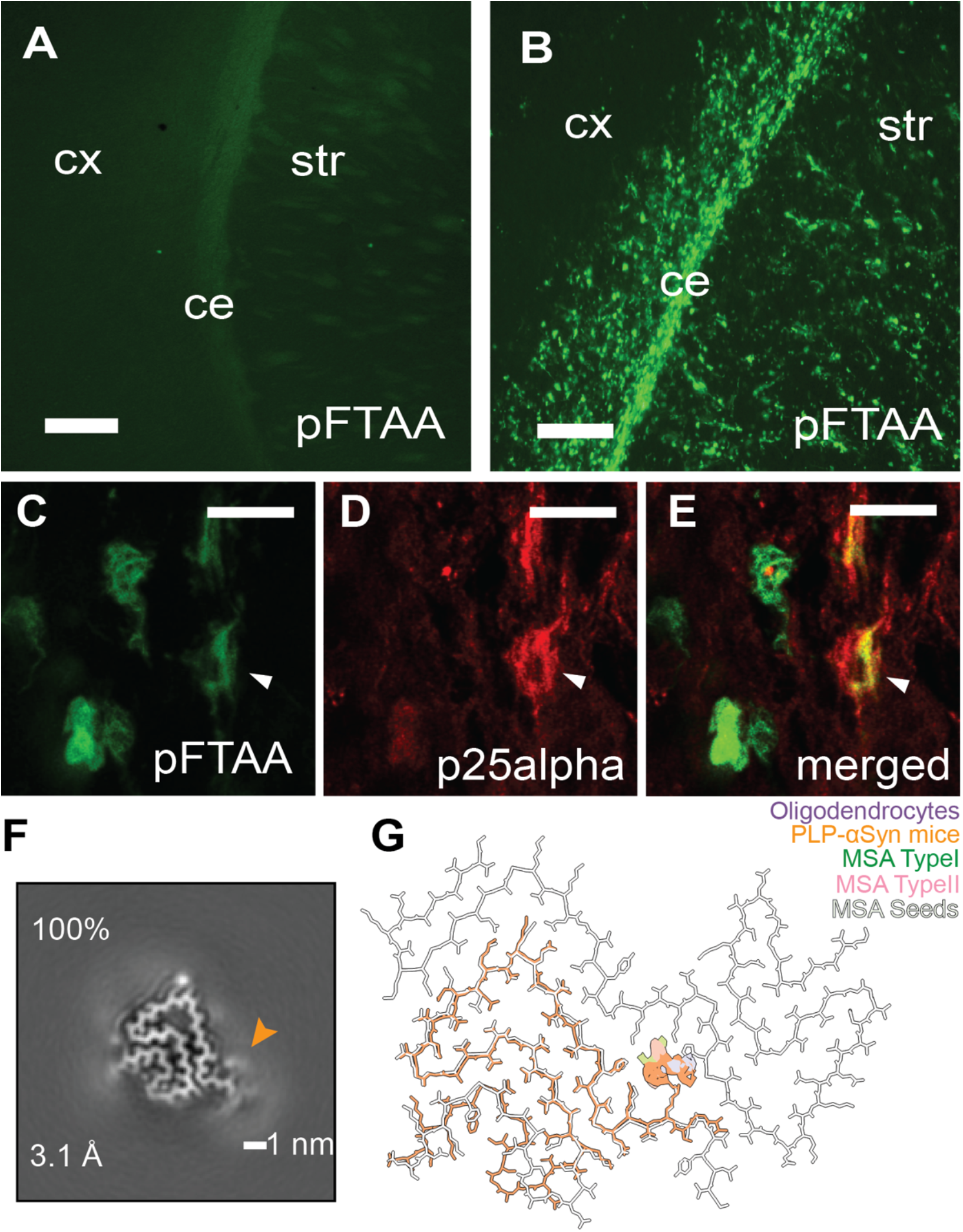
Intrastriatal injection of MSA seeds into PLP-αSyn transgenic mice induces the formation of amyloid inclusions in oligodendrocytes. A,B. pFTAA fluorescence in the brains of non-injected PLP-αSyn mice (A) and seven months following the intrastriatal injection of MSA seeds (B). C-E. Co-localisation of pFTAA staining and p25α immunoreactivity in the brains of PLP-αSyn mice injected with MSA seeds. Abbreviations: cx, cortex; ce, capsula externa; str, striatum. Scale bars, 200 μm (A, B); 10 μm (C-E). F. Cryo-EM cross-section of seeded α-synuclein filament from PLP-αSyn mice. G. Overlay of the atomic models of the seeded α-synuclein filament structure from PLP-αSyn mice with the model of the MSA seeds (1.9 Å resolution structure) and overlay of the non-proteinaceous densities in the cryo-EM maps of MSA-seeded α-synuclein filaments with those of α-synuclein filaments from the brains of individuals with MSA (MSA Type I, EMD-10650; MSA Type II_1_, EMD-45979).

Cryo-EM structure determination of the filaments again showed singlets with the PF-IIB_1_ conformation and an extra density between K43 and K45 (Figure 6F,G). The PF-IIB_1_ of the α-synuclein singlets was nearly identical to the corresponding structures obtained with the oligodendrocyte-enriched primary cultures described above with XY-APDs of 0 (Figure S1J). Unlike the seeded structures of α-synuclein filaments from oligodendrocyte-enriched primary cultures, there was another extra density connected to K60 in the seeded structures from PLP-αSyn transgenic mice.

## Discussion

MSA filaments represent an outlier among brain-derived amyloid filaments. Whereas many other amyloids also form doublets in the brain, for example tau, amyloid-β and TMEM106B, MSA filaments are the only ones known to have two protofilaments of the same protein in distinct conformations. The molecular drivers of this asymmetry remain unclear and none of the attempts to reconstitute MSA filaments in the laboratory have been successful. Seeded aggregation of recombinant α-synuclein did result in the formation of doublets in some cases^24,25^, but they were symmetrical and adopted filament structures different from those of MSA. The asymmetry of MSA filaments may be driven by distinct post-translational modifications of the α-synuclein molecules in the two protofilaments and such modifications may be provided by the cellular environment in which the filaments form.

*In vitro* cell cultures could provide the post-translational modifications that are required to break the symmetry in MSA filaments. However, our attempts at reconstituting MSA filaments by seeded aggregation in cell cultures were not successful. Whether using undifferentiated SH-SY5Y cells, human iPSC-derived neurons, mouse primary cultures or mouse-derived oligodendrocytes, we only ever observed singlet filaments. The observation that these singlets adopted the same conformation as one of the protofilaments of the MSA seeds indicates that templated seeding did occur, but it remains unclear why doublets did not form.

Possibly, the overexpression of α-synuclein, which we used to obtain enough seeded filaments for cryo-EM structure determination, is incompatible with doublet formation. Both transgenic mouse models that have been reported to yield singlets upon seeding with MSA filaments have also relied on overexpression: either human A53T α-synuclein in heterozygous M83 mice^48,49^ or human wildtype α-synuclein in homozygous PLP-αSyn mice^27^. Future experiments of seeded aggregation in cells and animals that do express physiological levels of α-synuclein may yield further insights. Such experiments would likely depend on large amounts of sample to allow cryo-EM structure determination. Another limitation is that we only used murine oligodendrocytes in this study. Human oligodendrocytes may be required for the formation of α-synuclein filaments made of two non-identical protofilaments.

In PLP-αSyn transgenic mice, we only detected amyloid filaments of α-synuclein in oligodendrocytes following the injection of MSA seeds. By contrast, pS129 α-synuclein staining was observed both without and with seed injection. It follows that pS129 staining is not a specific marker for α-synuclein filaments. Similar findings have been reported in PLP-αSyn mice following the intrastriatal injection of filaments of recombinant α-synuclein^50^.

The observation that seeded aggregation of MSA filaments in murine oligodendrocytes yielded a similar extra density between K43 and K45 as observed at the protofilament interface of MSA filaments, in both mouse primary cultures and in PLP-αSyn transgenic mice, suggests that oligodendrocytes provide the compounds that are responsible for this density in MSA filaments. Interestingly, the same singlets, but without the extra density between K43 and K45, have recently been described for seeded α-synuclein filaments from heterozygous M83 mice that had been injected with MSA seeds^48^. These mice overexpress human A53T α-synuclein downstream of the (largely neuronal) *Prnp* promoter and only develop abundant α-synuclein inclusions and clinical symptoms in the homozygous state^51^. However, when heterozygous M83 mice are injected with MSA seeds, they develop abundant neuronal α-synuclein inclusions and a progressive neurological disease with an incubation period of around 100 days^49^. Comparison of the seeded α-synuclein filament structures from mouse brains with those from the cellular model systems described above suggests that a neuronal environment alone is not sufficient for yielding MSA filaments and provides additional support for the hypothesis that the molecules responsible for the extra density at the protofilament interface of MSA filaments are provided by oligodendrocytes (Figure 7).

**Figure 7:**
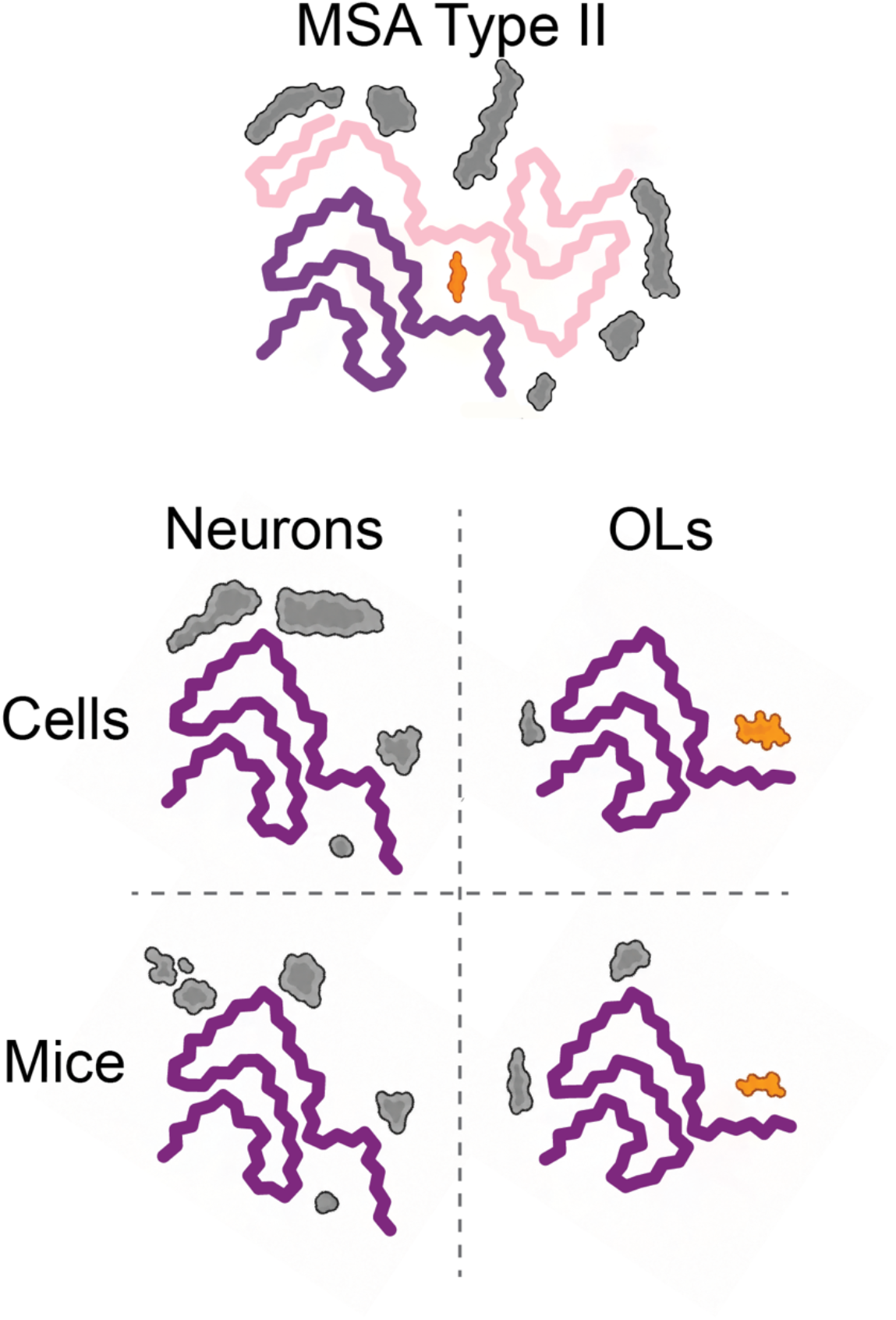
Summary. Seeding with Type II MSA filaments (top) led to singlets in cell cultures and mice overexpressing α-synuclein. Only overexpression in oligodendrocytes led to extra densities (orange) that were reminiscent of the extra density in MSA filaments, suggesting that oligodendrocytes may provide the molecules that make up those densities.

Whereas PLP-αSyn transgenic mice express high levels of human α-synuclein in oligodendrocytes, in the human brain, mature oligodendrocytes express α-synuclein at low levels compared to neurons^52,53^. In disease, the α-synuclein that ends up in MSA filaments is probably expressed in neurons and then imported into oligodendroglia. Whether at this point α-synuclein is already partially assembled or still monomeric is not known. It has been reported that oligodendrocytes can take up recombinant α-synuclein in a dynamin-dependent manner^54^. A primary role for oligodendrocytes is also supported by studies suggesting that the formation of GCIs may precede neuronal loss^55,56^. Our results indicate that singlets with the same conformation as that of one of the MSA protofilaments can form in both neurons and oligodendrocytes.

Although this work further implicates oligodendrocytes as key players in the formation of MSA filaments, it does not yet provide the full cellular model system needed to study their templated seeding. Such models have not been reported for any of the disease-specific α-synuclein filaments. It is our hope that continued efforts to form amyloid filaments with disease-specific structures will provide new insights into the molecular factors that drive the formation of these specific structures in diseased brains, and hence new avenues for therapeutic interventions.

## Materials and Methods

### MSA brains

Sarkosyl-insoluble extracts of the putamen from the previously reported MSA case 5 (PF-IIA/PF-IIB_1_ and PF-IIA/PF-IIB_2_ filaments)^19^ were used as seeds in the SH-SY5Y experiments. Extracts of the frontal cortex from the previously reported FTLD-synuclein case (PF-IIA/PF-IIB_1_ filaments)^35^ were used for Triton X-100 and sarkosyl extractions. Sarkosyl-insoluble extracts were used as seeds in human iNeurons, mixed primary murine cortical cells and primary murine oligodendrocytes, as well as for injection into PLP-αSyn transgenic mice.

### Sarkosyl extraction of MSA filaments and seed preparation

One gram of brain tissue was homogenised using a Polytron (PT 2500 E, Kinematica) in 20 vol (v/w) Buffer 1 (10 mM Tris-HCl, pH 7.4, 800 mM NaCl, 10% sucrose) supplemented with protease and protein phosphatase inhibitors (Sigma). Samples were adjusted to 2% sarkosyl and incubated at 220 rpm for 60 min at 37 °C. The homogenates were then centrifuged at 9,600 × g for 10 min at 25 °C, and the resulting supernatants centrifuged at 130,000 × g for 1 h at 25 °C. The pellets were resuspended in 1 ml of Buffer 1. The samples were sonicated with a Microson XI-2000 Ultrasonic Cell Disruptor (Misonix) for 20 s and diluted threefold with Buffer 2 (20 mM Tris-HCl, pH 7.4, 100 mM NaCl, 0.2% sarkosyl, 10% sucrose). All samples were centrifuged at 130,000 × g for 1 h at 25 °C. The pellets were resuspended in 500 µl PBS and centrifuged again at 130,000 × g for 1 h at 25 °C. The final pellets were resuspended in 50 µl Buffer 3 (20 mM Tris-HCl, pH 7.4, 100 mM NaCl) per gram of tissue.

For seeding experiments in iNeurons, mixed primary murine cortical cells and murine oligodendrocyte-enriched primary cultures, the final MSA sarkosyl-insoluble pellets were sonicated using a water-bath sonicator (Qsonica) at 50% amplitude, with cycles of 30 s on and 30 s off, for a total 4 min.

For seeding experiments in PLP-αSyn mice, the final MSA sarkosyl-insoluble pellets were suspended in 45 μl PBS and sonicated using a water-bath sonicator (Qsonica) at 50% amplitude, with cycles of 30 s on and 30 s off, for a total 6 min. The sonicated samples were then centrifuged at 9,000 x g for 2 min at 4 °C and the supernatants collected for subsequent injections.

### Triton X-100 extraction of MSA filaments

One gram frontal cortex from the FTLD-synuclein case^35^ was homogenised using a Polytron in 5 ml of Buffer 1 (10 mM Tris, pH 7.4, 800 mM NaCl, 10% sucrose), supplemented with protease and protein phosphatase inhibitors (Sigma). The samples were incubated at 4 °C for 20 min before centrifugation at 100,000 × g for 30 min at 4 °C. The pellets were then resuspended and homogenised using a Dounce homogeniser (Jencons) in 5 ml Buffer 4 (10 mM Tris, pH 7.4, 800 mM NaCl, 1% Triton X-100, 10% sucrose), supplemented with protease and protein phosphatase inhibitors (Sigma). The samples were incubated at 4 °C for 20 min before centrifugation at 100,000 × g for 30 min at 4 °C. The pellets were resuspended and homogenised again using a Dounce homogeniser in 5 ml Buffer 5 (10 mM Tris, pH 7.4, 800 mM NaCl, 1% Triton X-100, 30% sucrose), supplemented with protease and protein phosphatase inhibitors (Sigma) and incubated at 4 °C for 1 h. The samples were then centrifuged at 9,600 × g for 10 min at 25 °C. The supernatants were collected and mixed with 5 ml Buffer 6 (10 mM Tris, pH 7.4, 800 mM NaCl, 1% Triton X-100), to achieve a final sucrose concentration of 15% before centrifugation at 135,000 × g for 45 min at 25 °C. The pellets were resuspended in 500 µl Buffer 4 and centrifuged at 135,000 × g for 45 min at 25 °C. The final pellets were resuspended in 30 µl Buffer 3.

### Lentivirus production and purification

Lentivirus was constructed and produced as described previously^57^. Briefly, HEK293T Lenti-X cells (Takara) were plated in a T75 flask containing 12 ml DMEM medium (Gibco) supplemented with 10% foetal bovine serum (FBS) to achieve ∼50% confluency. The next day, the medium was replaced with 12 ml of DMEM (Gibco) supplemented with 2% FBS. Ten µg of the lentiviral transfer plasmid pHR-SFFV-WT-Venus or pHR-SFFV-WT-HA (adapted from Addgene), 10 µg of the envelope plasmid pMD2.G (Addgene) and 10 µg of the packaging plasmid psPAX2 (Addgene) were mixed with Lipofectamine 3,000 (Thermo Fisher Scientific) and incubated at room temperature for 30 min before addition to the cultures. After 48 h, the media were collected and filtered through a 0.45 µm filter (Millipore). Lenti-X Concentrator (Takara) was added to the filtered media at 4 °C for 2 h. The samples were then centrifuged at 1,500 × g for 45 min at 4 °C to pellet the viral particles. The final viral pellets from each T75 flask were resuspended in 170 µl of filtered PBS. The suspensions were divided into 20 µl aliquots and stored at −80 °C until use.

### Adeno-associated virus production and purification

Adeno-associated viruses (AAVs) were constructed and produced as described previously^58^. HEK293T AAVpro cells (Takara) were plated in a 15-cm dish containing 12 ml of DMEM supplemented with 10% FBS. When the cells had reached 80% confluence, the medium was replaced with penicillin/streptomycin-free medium. The following day, 7 µg of the AAV transfer plasmid carrying the transgene, 20 µg of the AAV helper plasmid and 7 µg of the AAV Rep/Cap plasmid were mixed with 1 mg/ml polyethylenimine (PEI) at a ratio of 5:1 and incubated at room temperature for 20 min before being added to the cultures. For mixed primary murine cortical cells, DNA encoding wildtype α-synuclein with a C-terminal Venus tag was cloned into an AAV transfer plasmid (Addgene) under the control of the human synapsin promoter. AAV Rep/Cap plasmids pAAV2/1 (Addgene) and pAAV2/2 (Addgene) were used for viral packaging. For oligodendrocyte-enriched primary cultures, DNA encoding wildtype α-synuclein with a C-terminal Venus tag was cloned into an AAV transfer plasmid (Addgene) under the control of the human myelin basic protein (MBP) promoter^44^. AAV Rep/Cap plasmid Olig001 (Addgene) was used for viral packaging. Following an overnight incubation, the medium was replaced with complete DMEM without FBS and the culture media were collected 48–60 h after transfection. The supernatants were incubated on ice with 8% PEG8000 and 318 mM NaCl for at least 2 h. The solutions were then centrifuged at 6,600 × g for 10 min to pellet the viral particles. The pellets were resuspended in 4 ml Buffer 7 (20 mM Tris, pH 8.0, 1 mM MgCl_2_, 150 mM NaCl). The cells were harvested and resuspended in 14 ml Buffer 7, combined with 4 ml virus particles, and subjected to three freeze–thaw cycles in the presence of Benzonase (Thermo Fisher Scientific) to release viral particles. The resulting solutions were centrifuged at 3,000 x g for 20 min at 4 °C. The supernatants were loaded onto an iodixanol (OptiPrep; Sigma) density gradient consisting of 17%, 25%, 40%, and 60% layers. After centrifugation at 340,000 x g for 70 min at 17 °C, the 40% fractions containing the viral particles were collected. They were then washed with PBS and concentrated using an Amicon 100K concentrator. The purity of the final viral preparations was assessed by SDS-PAGE, followed by aliquoting and storage at −80 °C until use.

### Human neuroblastoma SH-SY5Y cells

SH-SY5Y cells were maintained at 37 °C in a humidified incubator with 5% CO₂ in DMEM/F-12 medium (Sigma-Aldrich) supplemented with 10% foetal calf serum, penicillin-streptomycin-glutamine (Gibco), and MEM non-essential amino acids (Gibco). Cells were grown to 50%-60% confluence in collagen-coated 6-well plates and transfected with pcDNA3.1 encoding human wildtype, human A53T, or mouse wildtype α-synuclein (1.25 µg) using X-tremeGENE 9 (Roche Life Science), according to the manufacturer’s instructions. After 6-8 h, MSA seeds (1 µl) were introduced using MultiFectam (Promega) according to the manufacturer’s instructions. Cells were incubated for a further 3 days before analysis.

### Human iNeurons

NGN2-OPTi-OX-hPSCs (Bit Bio Ltd) were cultured in mTeSR base medium and maintained on plates coated with Cultrex basement membrane extract (R&D Systems) at 35 µg/cm², as described^40,59^. Cells were maintained at 37 °C in a humidified incubator with 5% CO₂ and fed every other day with mTeSR base medium. For iNeuron differentiation, stem cells were dissociated with Accutase (Thermo Fisher Scientific) and plated at a density of 0.6 x 10⁶ cells/ml in 6-well or 96-well plates coated with poly-D-lysine (50 µg/ml in borate buffer; Thermo Fisher Scientific) and Cultrex. After an overnight incubation, the medium was replaced with D0-2 induction medium consisting of DMEM/F-12 (Gibco), 1× GlutaMAX (Gibco), 1× non-essential amino acids (Gibco), 1× N2 supplement (Gibco), 1% penicillin-streptomycin, and doxycycline (1 µg/ml; Merck). After 48 h, cells were switched to D2 medium consisting of Neurobasal Plus (Gibco), 1× GlutaMAX, 1× B-27 Plus Supplement (Gibco), brain-derived neurotrophic factor (10 ng/ml; PeproTech), neurotrophin-3 (10 ng/ml; PeproTech), 1% penicillin-streptomycin, and doxycycline (1 µg/ml). Doxycycline treatment was maintained until day 7. Lentivirus encoding wildtype alpha-synuclein under spleen focus-forming virus (SFFV) promoter was added on day 4, followed by the addition of MSA seeds on day 6. Cells were harvested for further analysis on days 28-30.

### Mixed primary murine cortical cells

Mixed primary murine cortical cells were prepared from P0–P1 C57BL/6J-Ola mice, as described^60^. Cortical tissues were dissected, dissociated using trypsin and plated at a density of 1.0-1.2 x 10⁶ cells/well in 6-well plates or 30,000 cells per well in 96-well plates in Neurobasal Plus medium supplemented with 1× B-27 Plus Supplement, 10% horse serum, 1× GlutaMAX, and 1% penicillin-streptomycin. After cell attachment, the medium was replaced with Neurobasal Plus supplemented with 1× B-27 Plus Supplement, 1× GlutaMAX, and 1% penicillin-streptomycin. An adeno-associated virus (AAV) expressing wildtype α-synuclein fused to Venus under the control of the human synapsin-1 promoter was added on day 0. MSA seeds were added on day 4. Cells were harvested for further analysis on days 16-18.

### Murine oligodendrocyte-enriched primary cultures

OPCs were isolated from the cortices of P0–P1 C57BL/6J-Ola mice by sucrose density centrifugation^42^. Briefly, cortices from P0-P1 mice were dissected, dissociated in Accumax and digested with papain at 37 °C for 30 min. Cell suspensions were pelleted by centrifugation at 200 × g for 5 min and resuspended in medium containing 12% OptiPrep (Sigma). Following centrifugation at 200 × g for 15 min, the supernatants were diluted to 6% OptiPrep and centrifuged again at 200 × g for 15 min. Cells were then seeded into T75 flasks at a density of 1 × 10⁴ cells/cm² in OPC proliferation medium and cultured for 7 days. PDGF-AA (30 µg/mL) was added to the culture every other day. Upon reaching confluence, OPCs were washed with DPBS and dissociated with Accutase at 37 °C for 5 min. Dissociated OPCs were counted and combined with freshly isolated primary cortical cells at an OPC-to-primary-cell ratio of 2:1 to generate oligodendrocyte-enriched primary cultures. Cells were plated in co-culture medium 1, which was replaced with co-culture medium 2 after 4 h. On day 3, cultures were transduced with AAVs carrying the Olig001 capsid and expressing wildtype human α-synuclein fused to a C-terminal Venus tag under the control of the MBP promoter. MSA seeds were added on day 5. Cells were harvested for further analysis on days 28-31.

The OPC proliferation medium consisted of DMEM/F-12 supplemented with 1× GlutaMAX, 1× B-27 Supplement, PDGF-AA (30 µg/ml), bFGF (10 µg/ml), human epidermal growth factor (10 µg/ml), and 1% penicillin-streptomycin. Co-culture medium 1 consisted of Neurobasal Plus supplemented with 1× GlutaMAX, 1× B-27 Plus Supplement, 10% horse serum, and 1% penicillin-streptomycin. Co-culture medium 2 consisted of Neurobasal Plus supplemented with 1× GlutaMAX, 1× B-27 Plus Supplement, 40 ng/ml triiodothyronine (T3), 1× N2 Supplement, and 1% penicillin-streptomycin.

### PLP-αSyn transgenic mice and intrastriatal MSA seed injection

The transgenic mouse line expressing wildtype human α-synuclein under the control of the murine PLP promoter^27^ was bred and housed under temperature-controlled pathogen-free conditions with a 12 h light/dark cycle and free access to food and water. All *in vivo* experiments were performed according to the ARRIVE guidelines with the permission of the Austrian Federal Ministry of Science and Research (BMFWF-2022-0.429.387). Eleven mice at two months of age received a stereotaxic injection of MSA seeds (1.5 µl or 3 µl) in the left striatum under deep isoflurane inhalation anaesthesia using a 5 µl Hamilton syringe at coordinates relative to Bregma (mm): anterior-posterior +0.2 mm, medial-lateral +2.0 mm, and dorsal-ventral −3.0 mm at a rate of 0.1 µl per minute. The needle was left in place for at least 5 min after injection to minimise retrograde flow along the needle tract. Mice were placed on a heat-pad and monitored until complete recovery from anaesthesia. Seven months after seed injection, the mice were sacrificed using an overdose of thiopental (Medicamentum Pharma).

For filament extraction and analysis, the mice were cervically dislocated, their brains dissected and separated into hemispheres, followed by snap freezing in liquid nitrogen. For histopathology, mice were sacrificed by intracardial perfusion with 4% paraformaldehyde (PFA). Whole brains were post-fixed in 4% PFA at 4 °C overnight. Fixed tissues were cryoprotected in 30% sucrose, slowly frozen in 2-methylbutane (Sigma-Aldrich) and stored at −80 °C until further use. Free-floating 40 μm sections were cut using a cryostat (Leica Microsystems). For pFTAA staining, sections were washed with PBS and incubated for 30 min in 1.5 µM pFTAA (Sigma-Aldrich). For immunofluorescence, sections were washed several times in PBS and blocked for 1 h at room temperature with PBS containing 10% normal serum, 0.3% Triton X-100. The sections were then incubated with rabbit anti-pS129 α-synuclein ([EP1536Y] Abcam) or goat anti-TPPP/p25alpha (MyBioSource) antibody overnight at 4 °C with shaking. After three washes in PBS, they were incubated with Alexa-488-or Alexa-594-conjugated secondary antibodies (Invitrogen) for 1 h at room temperature. Following washes in PBS, the sections were embedded in fluorescence embedding medium (Invitrogen) and kept in the dark. Imaging was performed using a fluorescence microscope Leica DMI4000 and a confocal microscope Zeiss LSM 700.

### Filament extraction from seeded cells and seeded mouse brains

Transfected SH-SY5Y cells were washed with PBS and extracted with 1 ml of 1% sarkosyl in Buffer 8 (10 mM Tris, pH 7.5, 10% sucrose, 800 mM NaCl, 1 mM EGTA), followed by a 10 s pulse sonication with a Microson XI-2000 Ultrasonic Cell Disruptor (Misonix) setting at amplitude 2.5. After a 30 min incubation at 37 °C, cell extracts were ultracentrifuged at 113,000 × g for 20 min at 25 °C. The supernatants were removed and the pellets washed with 30 mM Tris-HCl (pH 7.5), followed by ultracentrifugation as before. The resulting pellets were the sarkosyl-insoluble fractions.

For seeded human iNeurons, mixed primary murine cortical cells and murine oligodendrocyte-enriched primary cultures, cells from one well of a 6-well plate were lysed in 500 µl of Buffer 9 (10 mM Tris, pH 7.4, 800 mM NaCl, 1% sarkosyl), supplemented with protease and protein phosphatase inhibitors (Sigma). The samples were then sonicated using a Microson XI-2000 Ultrasonic Cell Disruptor (Misonix) setting at amplitude 2.5 for 7 s and incubated at 37 °C with shaking at 600 rpm for 1 h. Lysates were centrifuged at 130,000 × g for 1 h at 25 °C. The pellets were washed with Buffer 10 (50 mM Tris, pH 7.4, 100 mM NaCl, 0.1% sarkosyl, 10% sucrose) and centrifuged again at 130,000 × g. Following washing, the pellets were resuspended in Buffer 3 (20 mM Tris, pH 7.4, 100 mM NaCl) for future experiments.

For seeded PLP-αSyn transgenic mice, the left hemispheres from three brains were pooled for sarkosyl extraction. Tissues were homogenised in 15 ml of Buffer 1 (10 mM Tris-HCl, pH 7.4, 800 mM NaCl, 10% sucrose), supplemented with protease and protein phosphatase inhibitors (Sigma). Samples were adjusted to 2% sarkosyl and incubated for 60 min at 37 °C while shaking. The homogenates were then centrifuged at 9,600 × g for 10 min at 25 °C, and the resulting supernatants centrifuged at 130,000 × g for 1 h at 25 °C. The pellets were resuspended in 250 µl of Buffer 1, diluted threefold with Buffer 2 (20 mM Tris-HCl, pH 7.4, 100 mM NaCl, 0.2% sarkosyl, 10% sucrose), followed by centrifugation at 130,000 × g for 1 h at 25 °C. The final pellets were suspended in Buffer 3 (20 mM Tris-HCl, pH 7.4, 100 mM NaCl) for future experiments. For non-injected PLP-αSyn mice, brainstems collected from six mice older than 12 months were used. The extraction procedures were the same as those used for injected PLP-αSyn mice.

### Immunohistochemistry

Cell cultures in 96-well plates were fixed with 4% paraformaldehyde, 4% sucrose, and 1% Triton X-100 in PBS or cold methanol, as previously described^61^. Following washes with PBS, cells were blocked with 2% BSA and 0.1% Triton X-100 in PBS for 1 h at room temperature and then incubated with primary antibodies overnight at 4 °C. Cells were washed three times with PBS and incubated with secondary antibodies for 2 h at room temperature. Following three additional washes with PBS, nuclei were stained with 1 µg/mL Hoechst 33342 (H3570) for 10 min at room temperature. Images were acquired at 488, 594 and 647 nm using a Nikon Ti2 inverted fluorescence microscope and analysed using Fiji software^62^.

### Immunogold electron microscopy and immunoblotting

Immunogold negative-stain electron microscopy and immunoblotting were carried out as described^63^. For immunogold negative-stain electron microscopy, 3 µl of appropriately diluted α-synuclein filament samples were applied to glow-discharged 400-mesh carbon film grids for 1 min. The grids were blocked with 0.1% gelatin in PBS for 5 min and incubated with anti-HA (Sigma, 1:50) or anti-GFP (Abcam, 1:50) antibodies for 1 h at room temperature. The grids were washed with 0.1% gelatin in PBS and incubated with a gold-conjugated secondary antibody (Sigma, 1:25) for 1 h. They were then washed and stained with 0.2% uranyl acetate. Images were acquired at 6,500× magnification using a Gatan Orius SC200B CCD detector on a Tecnai G2 Spirit operated at 120 kV.

For immunoblotting, samples were sonicated, mixed with loading buffer containing 2-mercaptoethanol and heated at 95 °C for 5 min before being loaded onto a Novex 4–12% Bis-Tris gel (Thermo Fisher Scientific). Proteins were transferred onto a nitrocellulose or a polyvinylidene fluoride (PVDF) membrane using a semi-dry transfer system (Bio-Rad). The membranes were fixed with 1% PFA and blocked with 2% BSA. The membranes were then incubated with antibodies Syn-1 (BD Biosciences, 1:2,000), pS129 (Abcam, 1:2,000), anti-HA (BioLegend, 1:4,000) or anti-GFP (Abcam, 1:4,000). Following this, the membranes were washed and incubated with the appropriate secondary antibodies: anti-mouse StarBright 700 (1:2,500) and anti-rabbit Alexa 800 (1:2,500). The membranes were then imaged using a ChemiDoc gel imager equipment (Bio-Rad).

### Electron cryo-microscopy

The extracted filaments were centrifuged at 3,000 g for 2 min to remove large debris. Holey carbon gold grids (Quantifoil Au R1.2/1.3, 300 mesh) were glow-discharged at 30 mA for 30 s using an Edwards E306A sputter coater. Aliquots of 3 µl were applied to the grids and blotted with Whatman filter paper at 100% humidity and 4 °C using a Vitrobot Mark IV (Thermo Fisher Scientific). Datasets for all samples were acquired using Titan Krios microscopes (Thermo Fisher Scientific) operated at 300 kV. Images of seeded filaments from SH-SY5Y cells overexpressing human α-synuclein, iNeurons overexpressing HA-tagged human α-synuclein, primary neurons overexpressing Venus-tagged human α-synuclein, PLP-αSyn transgenic mice overexpressing human α-synuclein and MSA-Triton X-100 samples were acquired using a Falcon 4i detector (Thermo Fisher Scientific). Images of seeded filaments from SH-SY5Y cells overexpressing human A53T α-synuclein or murine α-synuclein and from iNeurons expressing Venus-HA-tagged human α-synuclein were acquired using a K3 detector (Gatan) with a BioQuantum energy filter (Gatan) at a slit width of 20 eV. Images for the high-resolution MSA dataset were acquired using a Falcon 4i detector (Thermo Fisher Scientific) and a Selectris X energy filter (Thermo Fisher Scientific) at a slit width of 10 eV. The oligodendrocyte-seeded structures were calculated from a combination of images acquired using a Falcon 4i detector, a K3 detector with a BioQuantum energy filter at a slit width of 20 eV and a Falcon 4i detector with a Selectris X energy filter at a slit width of 10 eV. Images were recorded with a total electron dose of 40 electrons per Å².

### Helical reconstruction

Movie frames were gain-corrected, aligned, dose-weighted and summed using the motion correction program in RELION^64^. Contrast transfer function (CTF) parameters were estimated using CTFFIND-4.1^65^. All subsequent image-processing steps were performed using helical reconstruction methods in RELION-5^39^. α-Synuclein filaments were picked manually, except for those in the MSA_Triton dataset, which were picked by autopicker^36^. For all datasets, reference-free 2D classification was performed to select suitable segments for further processing. Initial 3D reference models were generated *de novo* from 2D class average images using relion_helix_inimodel2d^66^. Three-dimensional classification was performed to separate distinct conformations and remove suboptimal particles. Three-dimensional auto-refinements included optimisation of the helical twist and rise once resolutions reached 4.7 Å. Bayesian polishing^67^ and CTF refinement^68^ were used to improve resolution. Final reconstructions were sharpened using standard post-processing procedures in RELION, and final resolutions were estimated from Fourier shell correlations (FSCs) at 0.143 between independently refined half-maps^69^. The relion_helix_toolbox program was used to impose helical symmetry on the post-processed maps, which were then used for model building and refinement.

### Model building and refinement

Atomic models were built manually using Coot^70^ on the basis of published structures (PDB 9CX6). Model refinements were performed using Servalcat^71^, ISOLDE^72^ and REFMAC. Models were validated withy MolProbity^73^. Figures were prepared with ChimeraX^74^.

## Data availability

Cryo-EM maps were deposited to the EM Data Bank under the following accession codes: EMD-100438 for MSA_Triton; EMD-100441 for MSA_High_Res; EMD-100343 for SHY-WT-αSyn-I; EMD-100517 for SHY-WT-αSyn-II; EMD-100521 for SHY-A53T-αSyn-I; EMD-100434 for SHY-A53T-αSyn-II; EMD-100435 for SHY-Mouse-αSyn; EMD-100436 for iN-αSyn-HA; EMD-100444 for PN-Syn1-αSyn-Venus; EMD-100437 for Ol-MBP-αSyn-Venus-I; EMD-100440 for Ol-MBP-αSyn-Venus-II; EMD-100439 for MSA_PLP-αSyn_mice. Corresponding refined atomic models were deposited to the PDB under the following accession codes: 35EG for MSA_Triton; 35EK for MSA_High_Res; 35BV for SHY-WT-αSyn-I; 35GY for SHY-WT-αSyn-II; 35HC for SHY-A53T-αSyn-I; 35EC for SHY-A53T-αSyn-II; 35ED for SHY-Mouse-αSyn; 35EE for iN-αSyn-HA; 35EO for PN-Syn1-αSyn-Venus; 35EF for Ol-MBP-αSyn-Venus-I; 35EJ for Ol-MBP-αSyn-Venus-II; 35EH for MSA_PLP-αSyn_mice. Please address requests for materials to the corresponding authors. Source data are provided with this paper.

## Acknowledgements

We thank the families of the patients for donating brain tissues; T. Darling, J. Grimmett and I. Clayson for help with high-performance computing and the EM facility of the Medical Research Council (MRC) Laboratory of Molecular Biology for help with cryo-EM data acquisition. We acknowledge the Diamond Light Source for access and support of cryo-EM facilities at the UK’s Electron Bio-imaging Centre (under proposal BI37660-44). We thank Alex Fellow and Andrew Carter from the MRC-LMB for help with iNeuron cultures, and Thóra Káradóttir and Sebastian Timmler from the Cambridge Stem Cell Institute for help with oligodendrocyte cultures. We thank Gregory Merz for providing the cryo-EM map of seeded filaments from heterozygous M83 mice injected with seeds from MSA brains.

## Author contributions

P.C.C., K.H., Z.J., Masato Hasegawa and M.G. identified individuals with MSA. D.Z. prepared MSA seeds, performed seeding in human iPSC-derived neurons, mouse primary cultures and mouse-derived oligodendrocytes. M.M. and N.S. performed MSA seed injections in PLP-αSyn mice and carried out immunohistochemistry. A.T. and Masato Hasegawa performed MSA seeding in SH-SY5Y cells. D.Z., J.M. and T.K. performed lentiviral production and cultured iNeurons. D.Z. and Melissa Huang performed AAV purification and cultured murine cells. D.Z. collected the cryo-EM data, which were analysed by D.Z., A.G.M. and S.H.W.S. M.G. and S.H.W.S. supervised the project. All authors contributed to the writing of the manuscript.

## Copyright statement

For the purpose of open access, the MRC Laboratory of Molecular Biology has applied a CC BY public copyright licence to any Author Accepted Manuscript version arising.

## Declaration of generative AI and AI-assisted technologies in the manuscript preparation process

During the preparation of this work, the author(s) used ChatGPT for grammar checking, to analyse atomic models and to prepare parts of Figure 7. The author(s) reviewed and edited the output as needed and take full responsibility for the content of the published article.

---

**Supplementary Figure 1:**
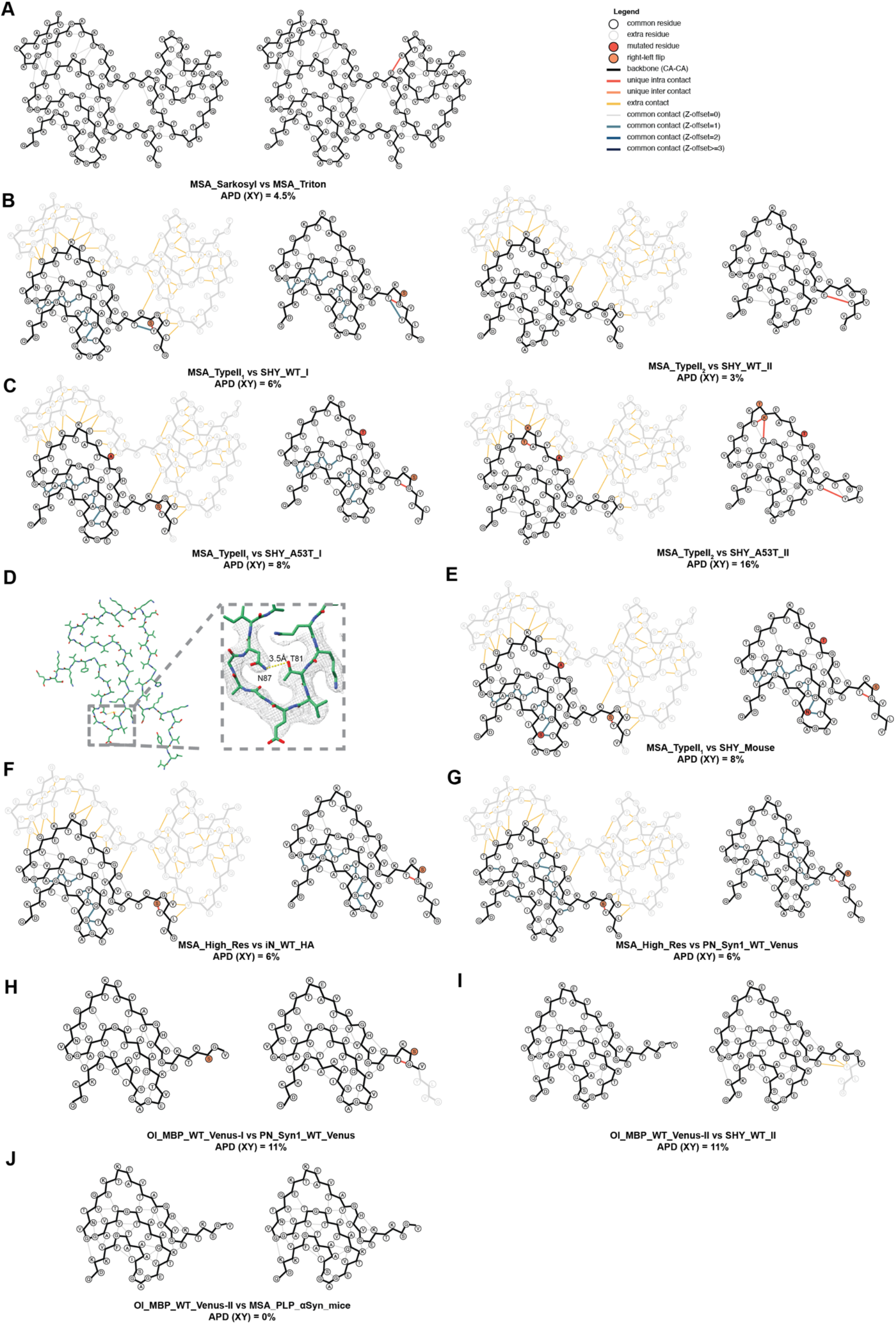
Amyloid packing differences. A. Amyloid packing difference (APD) between the structures of sarkosyl-extracted (PDB 9CX6) and Triton X-100-extracted filaments from the brains of individuals with MSA. B,C. APDs between the structures of MSA-seeded filaments from SH-SY5Y cells overexpressing wildtype or A53T human α-synuclein and those of the seeds (PDB 6XYP and 6XYQ). D. Potential hydrogen bond between N87 and T81 in the structures of MSA-seeded filaments from SH-SY5Y cells overexpressing murine wildtype α-synuclein. E. APD between the structures of MSA-seeded filaments from SH-SY5Y cells overexpressing wildtype murine α-synuclein and those of the seeds (PDB 6XYP). F,G. APDs between the 1.9 Å structure and the structure of MSA-seeded filaments from iNeurons overexpressing wildtype α-synuclein with a HA tag and from mixed primary murine cortical cells overexpressing wildtype α-synuclein with a Venus tag, respectively. H. APD between the type I structure of MSA-seeded filaments from murine oligodendrocyte-enriched primary cultures overexpressing wildtype α-synuclein with a Venus tag and the structure of MSA-seeded filaments from mixed primary murine cortical cells overexpressing wildtype α-synuclein with a Venus tag. I. APD between the type II structure of MSA-seeded filaments from murine oligodendrocyte-enriched primary cultures overexpressing wildtype α-synuclein with a Venus tag and the type II MSA-seeded α-synuclein filaments from SH-SY5Y cells overexpressing wildtype α-synuclein. J. APD between the structure of MSA-seeded filaments from PLP-αSyn mice and the type II structure of MSA-seeded filaments from murine oligodendrocyte-enriched primary cultures overexpressing wildtype α-synuclein with a Venus tag.

**Supplementary Figure 2:**
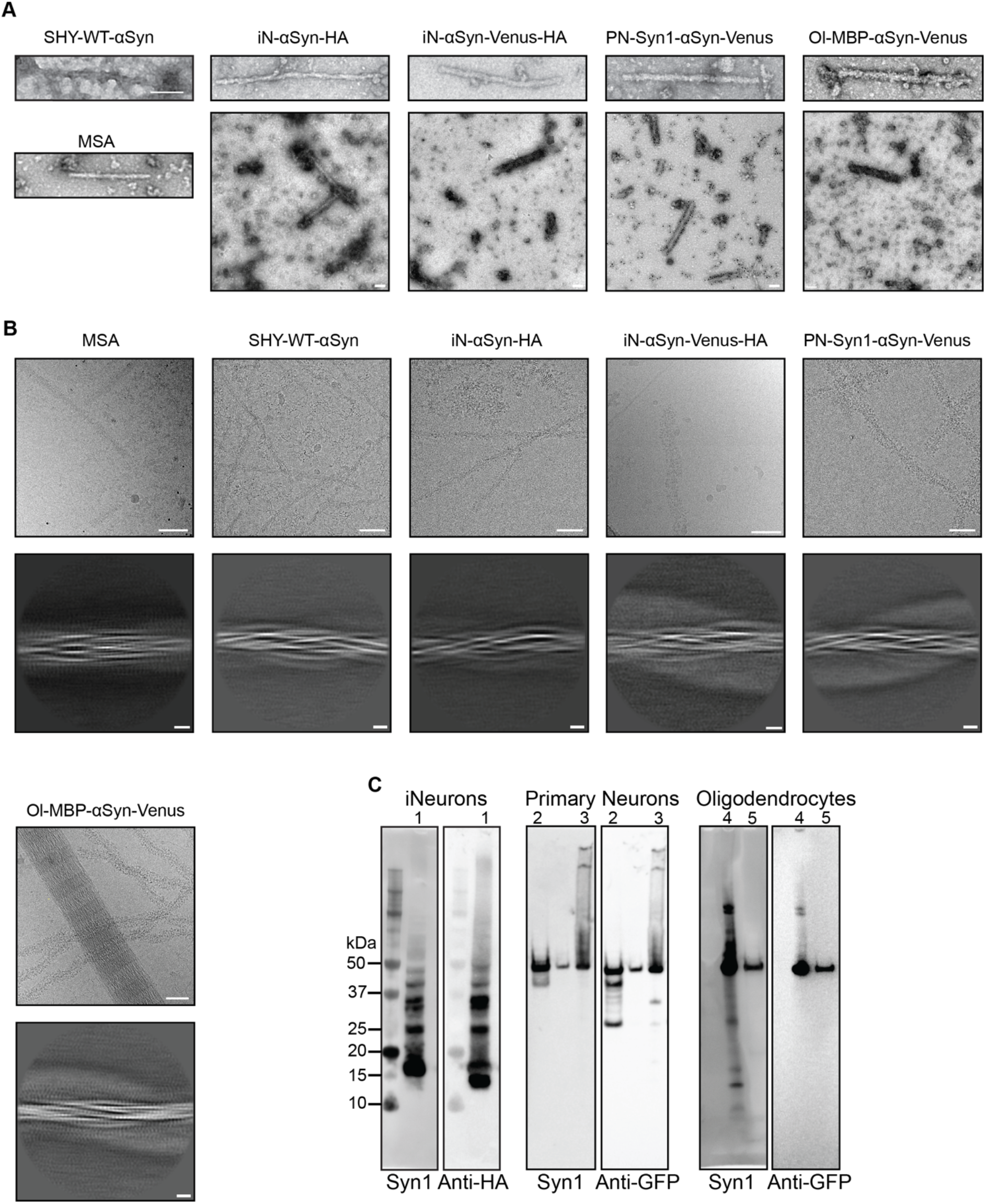
2D class averages of α-synuclein filaments with different tags (HA tag, Venus tag and Venus-HA tag) A. Negative-stain and immuno-EM images of α-synuclein filaments from the brains of individuals with MSA and MSA-seeded α-synuclein filaments from SH-SY5Y cells, iNeurons, mixed primary murine cortical cells and murine oligodendrocyte-enriched primary cultures. Scale bars, 100 nm. B. Cryo-EM images and 2D class averages of α-synuclein filaments from the brains of individuals with MSA and MSA-seeded α-synuclein filaments from SH-SY5Y cells, iNeurons, mixed primary murine cortical cells and murine oligodendrocyte-enriched primary cultures. Scale bars in cryo-EM images, 50 nm; scale bars in 2D class averages, 5 nm. C. Western blot analysis of α-synuclein filaments from MSA-seeded cells. Lanes: 1, sarkosyl-insoluble fractions from iNeurons; 2,3, sarkosyl-soluble (2) and sarkosyl-insoluble (3) fractions from mixed primary murine cortical cells; 4,5, sarkosyl-soluble (5) and sarkosyl-insoluble (4) fractions from murine oligodendrocyte-enriched primary cultures.

**Supplementary Figure 3:**
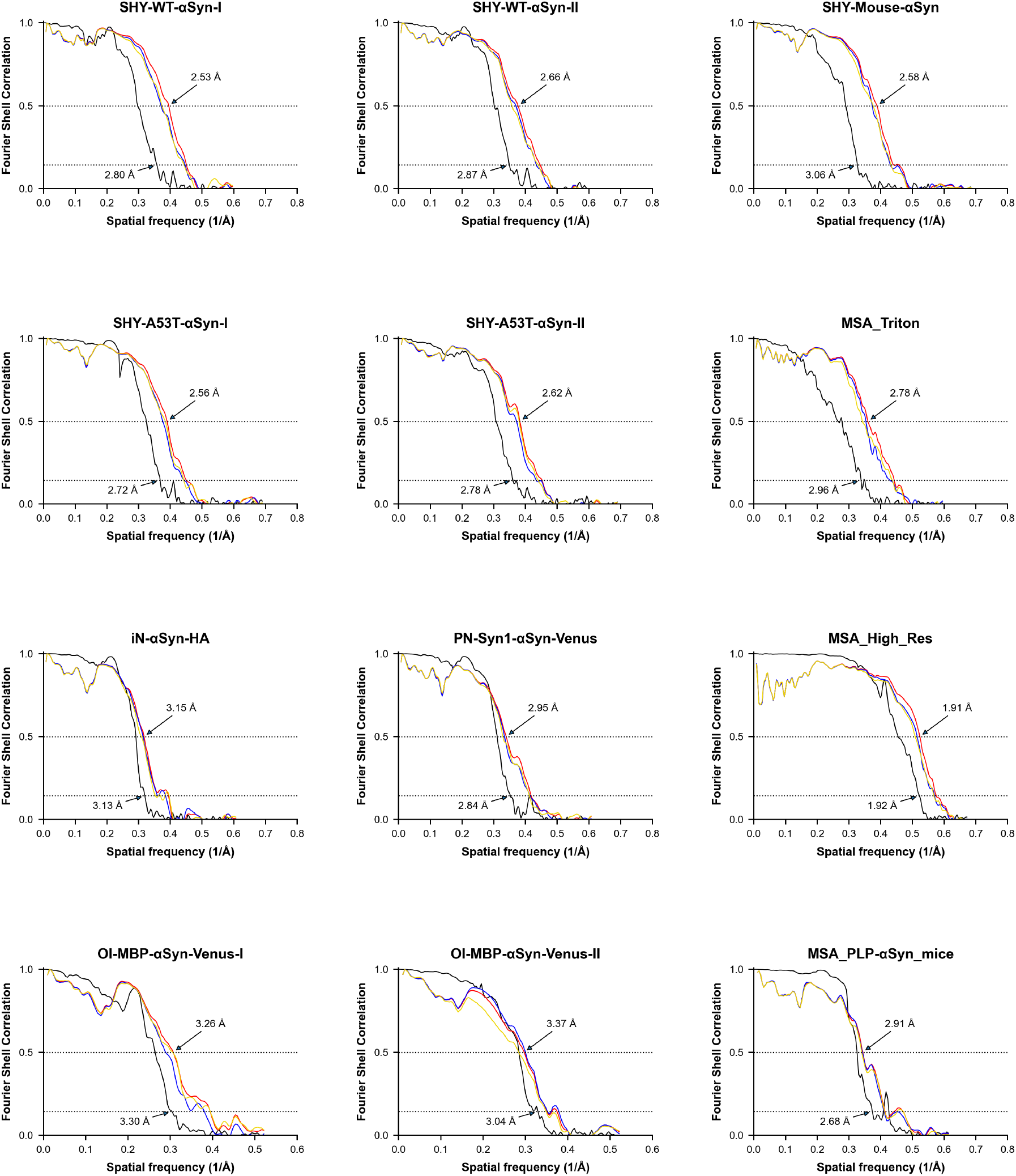
Fourier shell correlation (FSC) curves. FSC curves for cryo-EM maps are shown in black; for the final refined atomic model against the final cryo-EM map in red; for the atomic model refined in the first half map against the first half map in blue; for the refined atomic model in the first half map against the second half map in yellow.

**Supplementary Figure 4:**
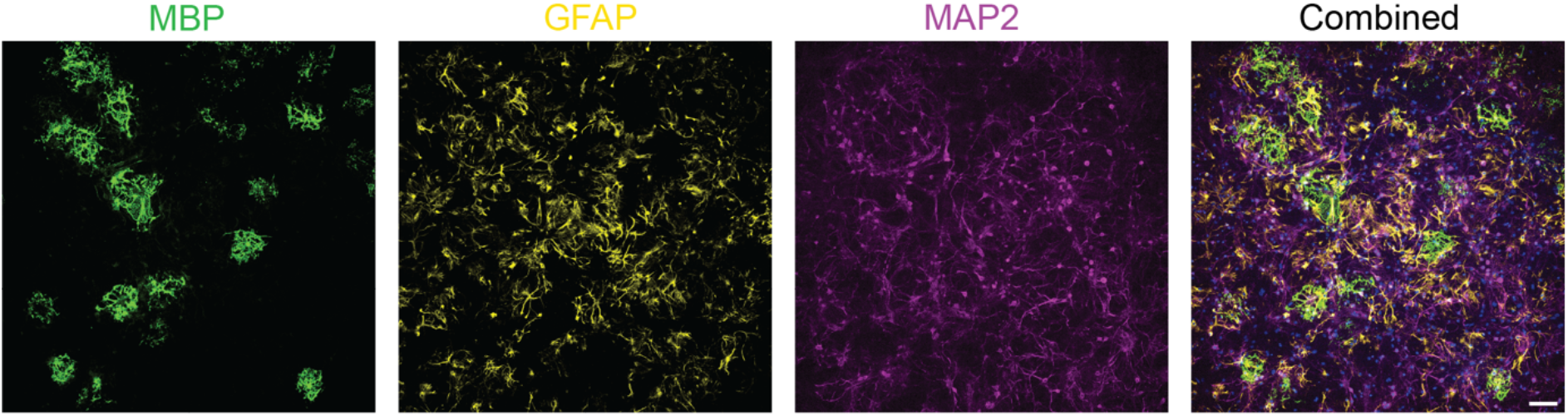
**Murine oligodendrocyte-enriched primary culture morphologies**. Markers show the presence of multiple cell types. MBP (green), GFAP (yellow), MAP2 (magenta), and DAPI (blue) identified oligodendrocytes, astrocytes, neurons and cell nuclei, respectively. Scale bar, 100 μm.

**Supplementary Figure 5:**
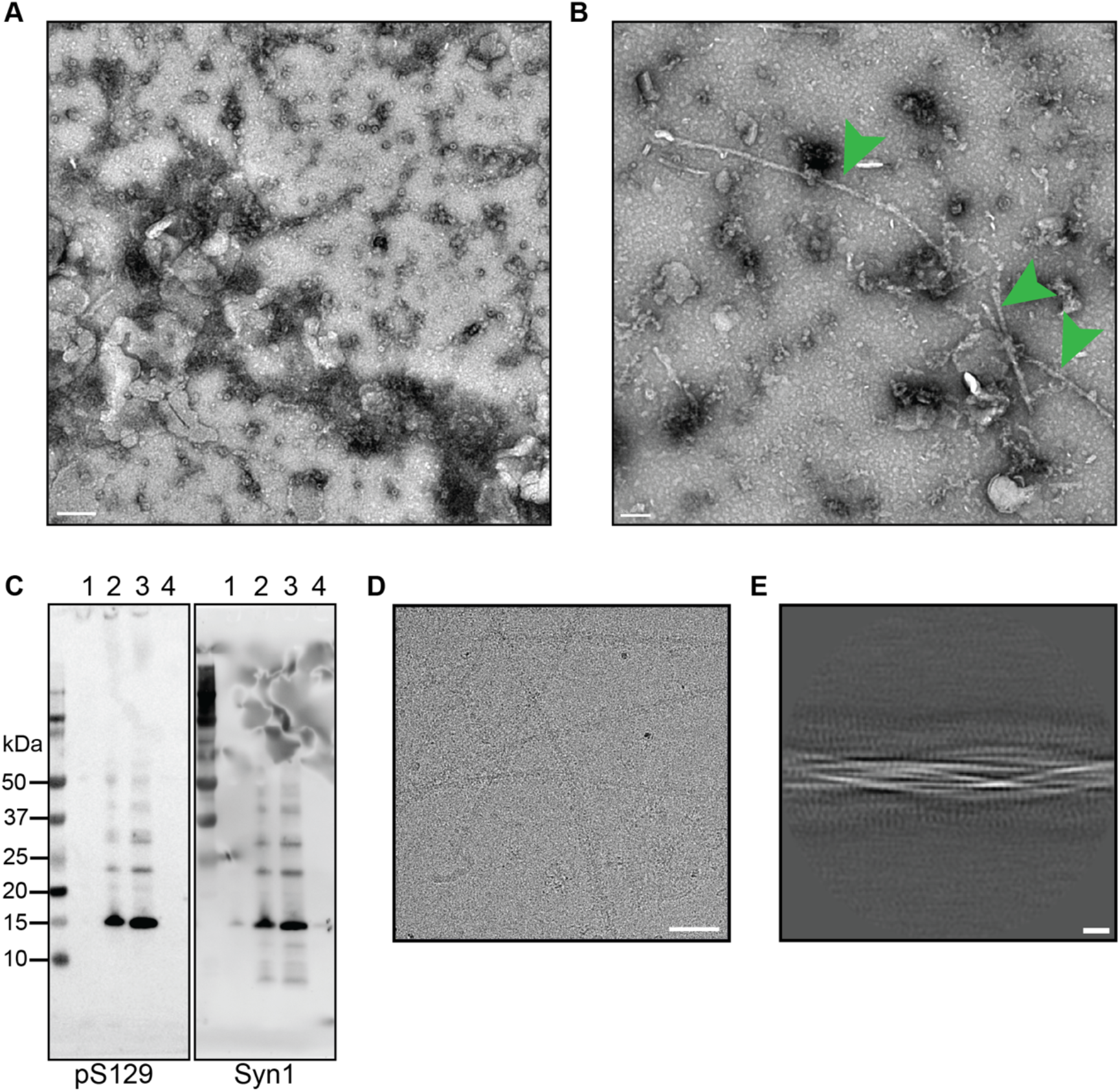
Intrastriatal injection of α-synuclein seeds from the brain of an individual with MSA induces filament assembly in PLP-αSyn mice. A. Negative-stain image of the sarkosyl-insoluble fraction from a non-injected PLP-αSyn mouse. No filaments are visible. Scale bar, 100 nm. B. Negative-stain image of the sarkosyl-insoluble fraction an MSA seed-injected PLP-αSyn mouse. Filaments are indicated by green arrows. Scale bar, 100 μm. C. Sarkosyl-insoluble fractions from the left (lane 1) and right (lane 4) hemispheres of non-injected PLP-αSyn mice, and from the left (lane 3) and right (lane 2) hemispheres of MSA seed-injected PLP-αSyn were probed with anti-α-synuclein antibodies pS129 and Syn1. D,E. Representative cryo-EM images (D) and 2D class averages (E) of filaments from MSA seed-injected PLP-αSyn mice. Scale bars: 50 nm (in D), 5 nm (in E).

**Supplementary Figure 6:**
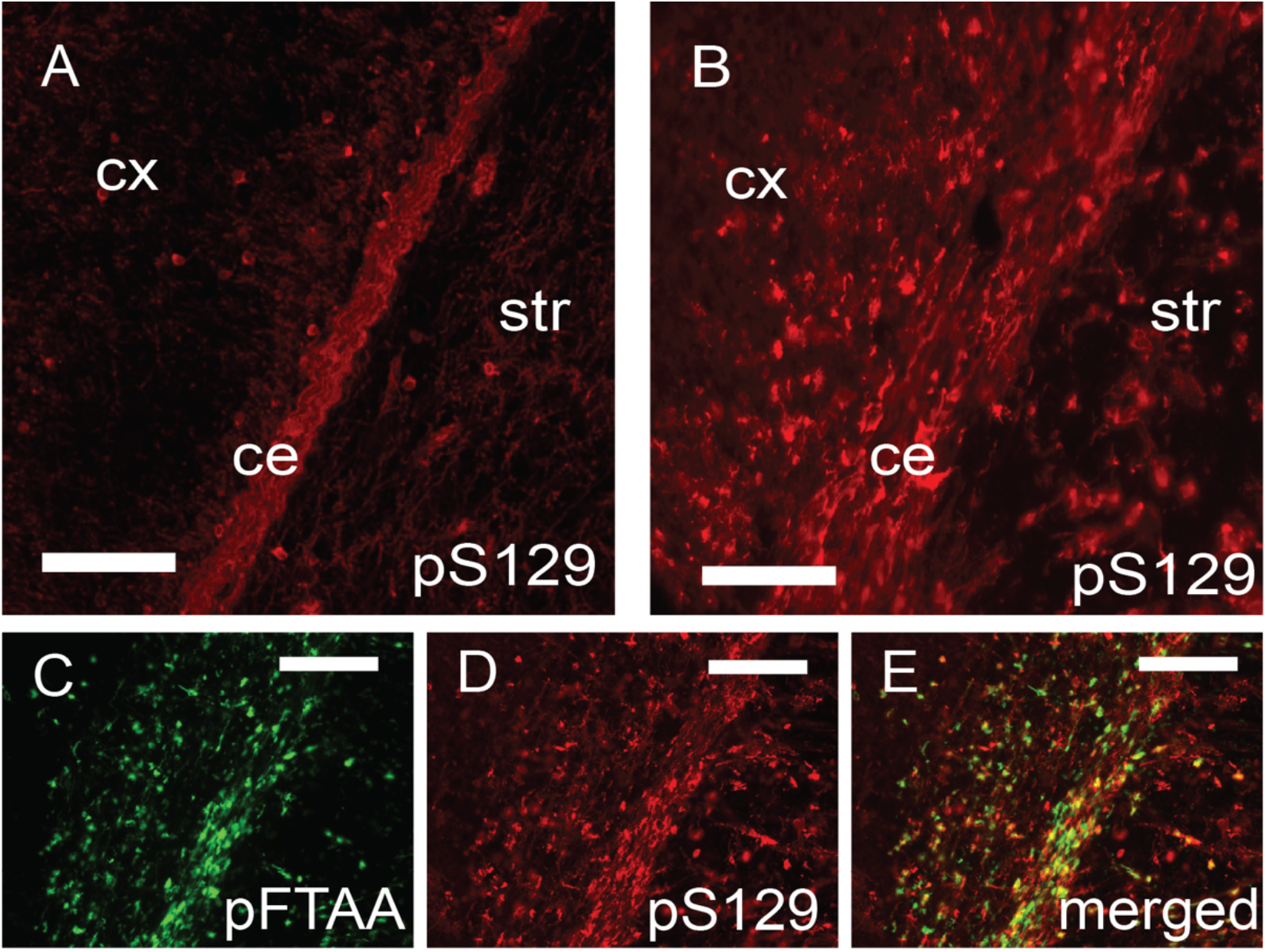
pS129 and pFTAA staining in non-injected and MSA seed-injected mice. A, B, Staining for α-synuclein phosphorylated at S129 in non-injected (A) and MSA seed-injected mice (B). C-E, Co-localisation of pFTAA staining and pS129 immunoreactivity in the brains of PLP-αSyn mice injected with MSA seeds. Scale bars, 200 μm (A, B); 180 μm (C-E).

**Supplementary Table 1:**
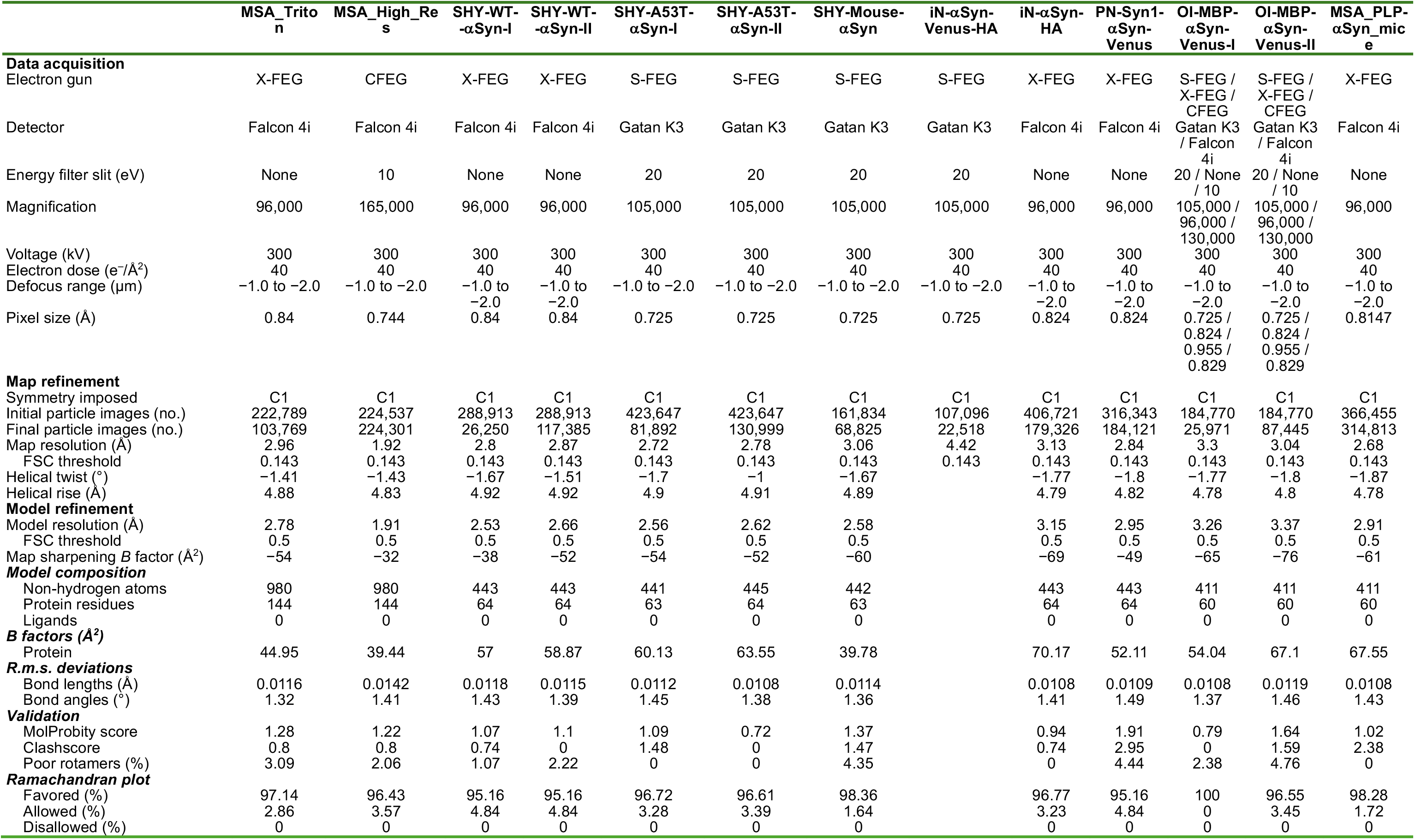
Cryo-EM data acquisition and structure determination.

|  | **MSA_Triton** | **MSA_High_Res** | **SHY-WT--αSyn-I** | **SHY-WT--αSyn-II** | **SHY-A53T-αSyn-I** | **SHY-A53T-αSyn-II** | **SHY-Mouse-αSyn** | **iN-αSyn-Venus-HA** | **iN-αSyn-HA** | **PN-Syn1-αSyn-Venus** | **OI-MBP-αSyn-Venus-I** | **OI-MBP-αSyn-Venus-II** | **MSA_PLP-αSyn_mice** |
|---|---|---|---|---|---|---|---|---|---|---|---|---|---|
| **Data acquisition** |  |  |  |  |  |  |  |  |  |  |  |  |  |
| Electron gun | X-FEG | CFEG | X-FEG | X-FEG | S-FEG | S-FEG | S-FEG | S-FEG | X-FEG | X-FEG | S-FEG / X-FEG / CFEG | S-FEG / X-FEG / CFEG | X-FEG |
| Detector | Falcon 4i | Falcon 4i | Falcon 4i | Falcon 4i | Gatan K3 | Gatan K3 | Gatan K3 | Gatan K3 | Falcon 4i | Falcon 4i | Gatan K3 / Falcon 4i | Gatan K3 / Falcon 4i | Falcon 4i |
| Energy filter slit (eV) | None | 10 | None | None | 20 | 20 | 20 | 20 | None | None | 20 / None / 10 | 20 / None / 10 | None |
| Magnification | 96,000 | 165,000 | 96,000 | 96,000 | 105,000 | 105,000 | 105,000 | 105,000 | 96,000 | 96,000 | 105,000 / 96,000 / 130,000 | 105,000 / 96,000 / 130,000 | 96,000 |
| Voltage (kV) | 300 | 300 | 300 | 300 | 300 | 300 | 300 | 300 | 300 | 300 | 300 | 300 | 300 |
| Electron dose (e<sup>−</sup>/Å<sup>2</sup>) | 40 | 40 | 40 | 40 | 40 | 40 | 40 | 40 | 40 | 40 | 40 | 40 | 40 |
| Defocus range (µm) | −1.0 to −2.0 | −1.0 to −2.0 | −1.0 to −2.0 | −1.0 to −2.0 | −1.0 to −2.0 | −1.0 to −2.0 | −1.0 to −2.0 | −1.0 to −2.0 | −1.0 to −2.0 | −1.0 to −2.0 | −1.0 to −2.0 | −1.0 to −2.0 | −1.0 to −2.0 |
| Pixel size (Å) | 0.84 | 0.744 | 0.84 | 0.84 | 0.725 | 0.725 | 0.725 | 0.725 | 0.824 | 0.824 | 0.725 / 0.824 / 0.955 / 0.829 | 0.725 / 0.824 / 0.955 / 0.829 | 0.8147 |
| **Map refinement** |  |  |  |  |  |  |  |  |  |  |  |  |  |
| Symmetry imposed | C1 | C1 | C1 | C1 | C1 | C1 | C1 | C1 | C1 | C1 | C1 | C1 | C1 |
| Initial particle images (no.) | 222,789 | 224,537 | 288,913 | 288,913 | 423,647 | 423,647 | 161,834 | 107,096 | 406,721 | 316,343 | 184,770 | 184,770 | 366,455 |
| Final particle images (no.) | 103,769 | 224,301 | 26,250 | 117,385 | 81,892 | 130,999 | 68,825 | 22,518 | 179,326 | 184,121 | 25,971 | 87,445 | 314,813 |
| Map resolution (Å) | 2.96 | 1.92 | 2.8 | 2.87 | 2.72 | 2.78 | 3.06 | 4.42 | 3.13 | 2.84 | 3.3 | 3.04 | 2.68 |
| FSC threshold | 0.143 | 0.143 | 0.143 | 0.143 | 0.143 | 0.143 | 0.143 | 0.143 | 0.143 | 0.143 | 0.143 | 0.143 | 0.143 |
| Helical twist (°) | −1.41 | −1.43 | −1.67 | −1.51 | −1.7 | −1 | −1.67 |  | −1.77 | −1.8 | −1.77 | −1.8 | −1.87 |
| Helical rise (Å) | 4.88 | 4.83 | 4.92 | 4.92 | 4.9 | 4.91 | 4.89 |  | 4.79 | 4.82 | 4.78 | 4.8 | 4.78 |
| **Model refinement** |  |  |  |  |  |  |  |  |  |  |  |  |  |
| Model resolution (Å) | 2.78 | 1.91 | 2.53 | 2.66 | 2.56 | 2.62 | 2.58 |  | 3.15 | 2.95 | 3.26 | 3.37 | 2.91 |
| FSC threshold | 0.5 | 0.5 | 0.5 | 0.5 | 0.5 | 0.5 | 0.5 |  | 0.5 | 0.5 | 0.5 | 0.5 | 0.5 |
| Map sharpening *B* factor (Å<sup>2</sup>) | −54 | −32 | −38 | −52 | −54 | −52 | −60 |  | −69 | −49 | −65 | −76 | −61 |
| ***Model composition*** |  |  |  |  |  |  |  |  |  |  |  |  |  |
| Non-hydrogen atoms | 980 | 980 | 443 | 443 | 441 | 445 | 442 |  | 443 | 443 | 411 | 411 | 411 |
| Protein residues | 144 | 144 | 64 | 64 | 63 | 64 | 63 |  | 64 | 64 | 60 | 60 | 60 |
| Ligands | 0 | 0 | 0 | 0 | 0 | 0 | 0 |  | 0 | 0 | 0 | 0 | 0 |
| ***B factors (Å<sup>2</sup>)*** |  |  |  |  |  |  |  |  |  |  |  |  |  |
| Protein | 44.95 | 39.44 | 57 | 58.87 | 60.13 | 63.55 | 39.78 |  | 70.17 | 52.11 | 54.04 | 67.1 | 67.55 |
| ***R.m.s. deviations*** |  |  |  |  |  |  |  |  |  |  |  |  |  |
| Bond lengths (Å) | 0.0116 | 0.0142 | 0.0118 | 0.0115 | 0.0112 | 0.0108 | 0.0114 |  | 0.0108 | 0.0109 | 0.0108 | 0.0119 | 0.0108 |
| Bond angles (°) | 1.32 | 1.41 | 1.43 | 1.39 | 1.45 | 1.38 | 1.36 |  | 1.41 | 1.49 | 1.37 | 1.46 | 1.43 |
| ***Validation*** |  |  |  |  |  |  |  |  |  |  |  |  |  |
| MolProbity score | 1.28 | 1.22 | 1.07 | 1.1 | 1.09 | 0.72 | 1.37 |  | 0.94 | 1.91 | 0.79 | 1.64 | 1.02 |
| Clashscore | 0.8 | 0.8 | 0.74 | 0 | 1.48 | 0 | 1.47 |  | 0.74 | 2.95 | 0 | 1.59 | 2.38 |
| Poor rotamers (%) | 3.09 | 2.06 | 1.07 | 2.22 | 0 | 0 | 4.35 |  | 0 | 4.44 | 2.38 | 4.76 | 0 |
| ***Ramachandran plot*** |  |  |  |  |  |  |  |  |  |  |  |  |  |
| Favored (%) | 97.14 | 96.43 | 95.16 | 95.16 | 96.72 | 96.61 | 98.36 |  | 96.77 | 95.16 | 100 | 96.55 | 98.28 |
| Allowed (%) | 2.86 | 3.57 | 4.84 | 4.84 | 3.28 | 3.39 | 1.64 |  | 3.23 | 4.84 | 0 | 3.45 | 1.72 |
| Disallowed (%) | 0 | 0 | 0 | 0 | 0 | 0 | 0 |  | 0 | 0 | 0 | 0 | 0 |

